# Neural manifolds in V1 change with top-down signals from V4 targeting the foveal region

**DOI:** 10.1101/2023.06.14.544966

**Authors:** Aitor Morales-Gregorio, Anno C. Kurth, Junji Ito, Alexander Kleinjohann, Frédéric V. Barthélemy, Thomas Brochier, Sonja Grün, Sacha J. van Albada

**Affiliations:** Institute for Advanced Simulation (IAS-6), Jülich Research Centre, Jülich, Germany; Institute of Zoology, University of Cologne, Cologne, Germany; RWTH Aachen University, Aachen, Germany; Theoretical Systems Neurobiology, RWTH Aachen University, Aachen, Germany; Institut de Neurosciences de la Timone (INT), CNRS & Aix-Marseille Université, Marseille, France; JARA-Institut Brain Structure-Function Relationships (INM-10), Jülich Research Centre, Jülich, Germany

**Keywords:** Neural manifold, dimensionality, electrophysiology, macaque, visual cortex, top-down signals, resting state

## Abstract

High-dimensional brain activity is often organized into lower-dimensional neural manifolds. However, the neural manifolds of the visual cortex remain understudied. Here, we study large-scale multielectrode electrophysiological recordings of macaque (*Macaca mulatta*) areas V1, V4 and DP with a high spatio-temporal resolution. We find, for the first time, that the population activity of V1 contains two separate neural manifolds, which correlate strongly with eye closure (eyes open/closed) and have distinct dimensionalities. Moreover, we find strong top-down signals from V4 to V1, particularly to the foveal region of V1, which are significantly stronger during the eyes-open periods, a previously unknown effect. Finally, *in silico* simulations of a balanced spiking neuron network qualitatively reproduce the experimental findings. Taken together, our analyses and simulations suggest that top-down signals modulate the population activity of V1, causing two distinct neural manifolds. We postulate that the top-down modulation during the eyes-open periods prepares V1 for fast and efficient visual responses, resulting in a type of visual stand-by state.

## Introduction

The brain can be described as a high-dimensional dynamical system capable of representing and processing a plethora of low-dimensional variables.

The time-resolved activity of a population of neurons can be considered as a trajectory in a high-dimensional space, where each neuron represents one dimension; i.e., the state space of the neural system. Typically, the system does not attain all possible states in the state space, but rather remains confined to small subsets. These subsets of the state space are referred to as a neural manifolds ^1–5^. Neural manifolds have been shown to encode aspects such as decision-making in the prefrontal cortex of macaque^6^, hand movement trajectories in the motor cortex of macaque^2,3,7^, odor in the piriform cortex of mice ^8^, head direction in the anterodorsal thalamic nucleus of mice^5^, and spatial position in the hippocampus of mice ^9^. The study of neural manifolds in the visual cortex has been conducted in mice ^10,11^, macaque population ^12^, and macaque complex single neuron ^13^ visual responses. However, to the best of our knowledge, the state dependence of neural manifolds in the primary visual cortex (V1) of macaque has not yet been investigated.

Neural manifolds often have an intricate structure, which can be studied using methods borrowed from computational topology^5,12,14^. In addition to the topology, the number of uncorrelated covariates required to capture the variance in the state space is studied as a measure of the dimensionality of a neural system^1,10,15–20^. Regardless of species and brain area, the dimensionality is drastically lower than the total number of recorded neurons (i.e., state-space dimension)^1^, suggesting robust encoding of low-dimensional variables. Stringer et al. ^10^ showed that the dimensionality of visual cortical activity in mice can vary dynamically to encode precise visual input, seen as changes in the power law exponent of the explained variance. Such dynamical changes in dimensionality have not yet been demonstrated in other species.

Whether a subject has its eyes open or closed is known to affect the activity in the visual cortex, even in darkness^21–25^. In particular, the spectral power in the alpha frequency band (roughly 8–12 Hz) is known to decrease when the eyes are open, commonly known as alpha blocking^26–28^. Alpha blocking is usually attributed to desynchronisation ^27^ or oscillatory damping^28^ within V1. However, the concrete pathway(s) triggering these phenomena, and the relation between eye closure and neural manifolds in V1, are still unknown.

The primary visual cortex (V1) is known to represent fine details of visual input at both single-neuron and population levels ^12,29^. The visual system is hierarchical in nature, with information travelling from lower to higher areas (bottom-up) and vice versa (top-down), within specific frequency bands^30–33^. Top-down signals from V4 to V1 are known to mediate visual attention for figure-ground segregation and contour integration in macaque^33–37^. Recent evidence suggests that top-down signals can modulate neural manifold geometry and their dimensionality^38,39^. Naumann et al. ^38^ show *in silico* that top-down signals can rotate neural manifolds to maintain context-invariant representations. Dahmen et al.^39^ show that recurrent connectivity motifs modulate the dimensionality of the cortical activity. As effective connectivity is input-dependent^40^, a change in top-down input may therefore affect the dimensionality of neural activity. However, whether top-down signals modulate neural manifold geometry and dimensionality *in vivo* remains to be shown.

Here, we study the state space of the primary visual cortex of the macaque (N=3) during the resting state and its relation to the top-down signals from higher visual areas (V4, DP). We find that the population activity of macaque V1 is organized as two distinct high-dimensional neural manifolds, which are correlated with the behavior (eye closure) of the macaques, but not related to any visual stimuli. The dimensionality of each of these manifolds is significantly different, with higher dimensionality found during the eyes-open periods than the eyes-closed periods. In addition, we estimated input from higher cortical areas to V1 and found that these top-down signals are significantly stronger during the eyes-open periods, suggesting they play a role in modulating the neural manifolds and their dimensionality. Finally, we simulate a spiking neuron model under resting-state conditions and show that top-down signals can induce multiple manifolds by changing the firing modes of the network. Taken together, the data analysis and simulations show that top-down signals could actively modulate the V1 population activity, leading to two distinct neural manifolds of macaque visual cortical activity.

## Results

To explore the activity in the visual cortex, the intracortical electrical potential from the visual cortex of three rhesus macaques (*Macaca mulatta*) was recorded. The experiments simultaneously recorded the activity from V1 and V4 (macaques L & A)^42^ and from V1 and DP (macaque Y, see Figure 1b)^43^. The recordings were made in the resting state, i.e., the macaques sat head-fixed in a dark room and were not instructed to perform any particular task. In this state the macaques often showed signs of sleepiness and kept their eyes closed for periods of variable duration. The right eye—contralateral to the site of neural recording—was tracked using an infrared camera, allowing the identification of periods of open or closed eyes. See methods Electrophysiological data from macaques L & A and Electrophysiological data from macaque Y for further details on the data acquisition and processing. The experimental setup and data processing steps are illustrated in Figure 1.

**Figure 1:**
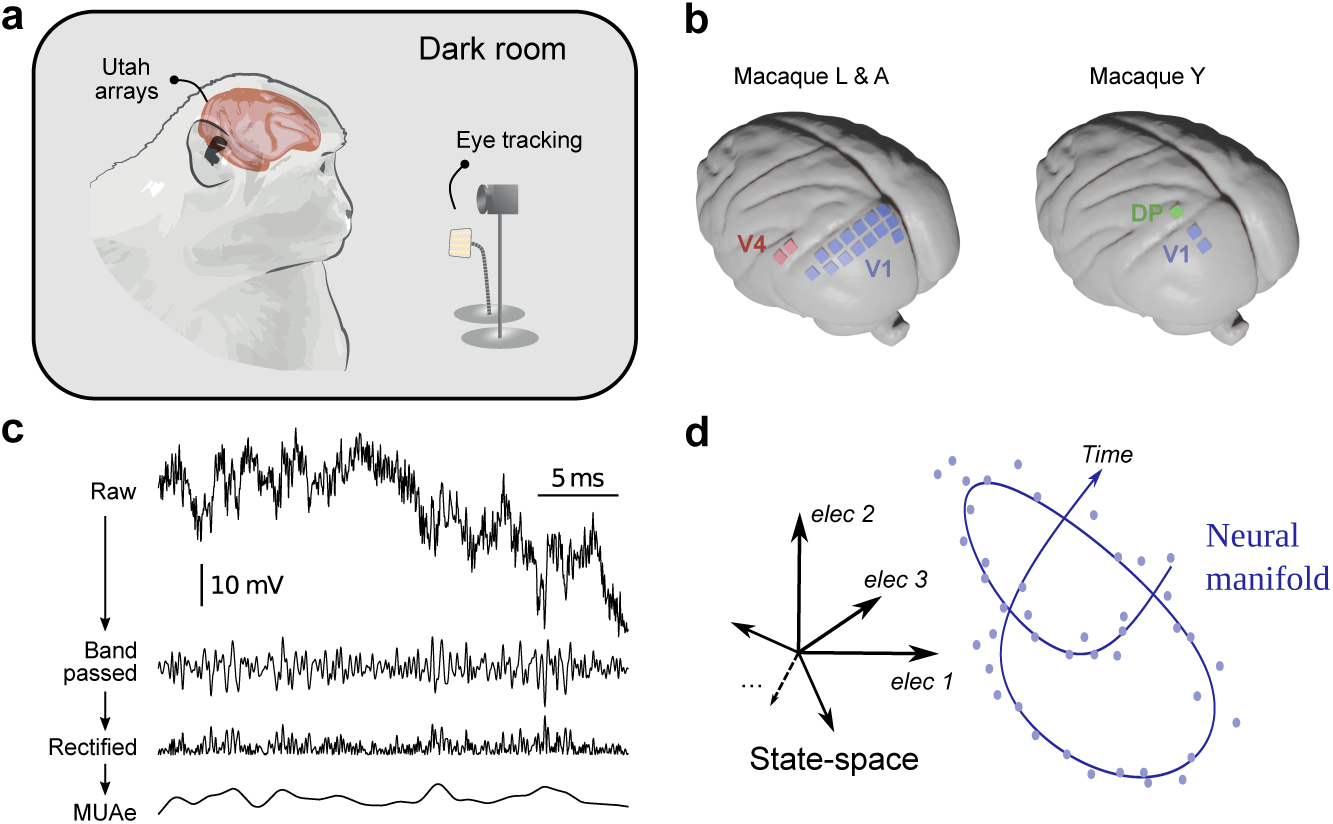
Overview of the experiment and neural manifold construction. **a** Illustration of the experimental setup. **b** Approximate locations of array implants in both experiments. Exact placement of the arrays differs slightly between subjects L and A. **c** Steps for obtaining the multi-unit activity envelope (MUAe) ^41^, used in this study. Band-pass filtering is performed between 500 Hz and 9 kHz, and the rectified signal is low-passed at 200 Hz to obtain the MUAe. **d** Schematic representation of state space and a neural manifold. Note that time is implicit within the neural manifold.

### Two distinct neural manifolds in V1 correlated with eye closure

To explore the activity of the visual cortex, we characterize the high-dimensional population activity (between 40 and 800 electrodes, see Table 1 for details) for each area and macaque in terms of the downsampled (1 Hz) multi-unit activity envelope (MUAe)^41^ (Figure 2a). We projected the population activity into a 3D space for visualisation using principal component analysis (PCA) (Figure 2b-d).

**Table 1:** Summary of subjects and recordings included in this study.

| Subject | Session | Duration (s) | Areas | Electrodes | Clean Electrodes |
| --- | --- | --- | --- | --- | --- |
| L | L_RS_250717 | 1363 | V1 | 896 | 765 |
|  |  |  | V4 | 128 | 116 |
| L | L_RS_090817 | 1321 | V1 | 896 | 761 |
|  |  |  | V4 | 128 | 116 |
| L | L_RS_100817 | 1298 | V1 | 896 | 774 |
|  |  |  | V4 | 128 | 118 |
| A | A_RS_150819 | 2278 | V1 | 896 | 402 |
|  |  |  | V4 | 128 | 11 |
| A | A_RS_160819 | 2441 | V1 | 896 | 369 |
|  |  |  | V4 | 128 | 9 |
| Y | Y_RS_180122 | 906 | V1 | 72 | 42 |
|  |  |  | DP | 36 | 25 |
| Y | Y_RS_180201 | 699 | V1 | 72 | 44 |
|  |  |  | DP | 36 | 24 |

**Table 2:** Peak detection algorithm parameters.

**CWT peak detection parameters**
|  |  |  |
| --- | --- | --- |
| widths | 100 – 500 Hz | Width range for CWT matrix. |
| width_step | 0.1 Hz | Step between widths. |
| wavelet | Ricker | Wavelet used for convolution. |
| max_distances | widths / 4 | Criterion to consider ridge lines connected. |
| gap_thresh | 10 Hz | Ridge lines farther apart will not be connected. |
| min_length | 225 | Minimum length of ridge lines. |
| min_snr | 1 | Minimum snr of ridge lines. |
| noise_perc | 10 | Percentile of ridge line considered noise for snr calculation. |

**Figure 2:**
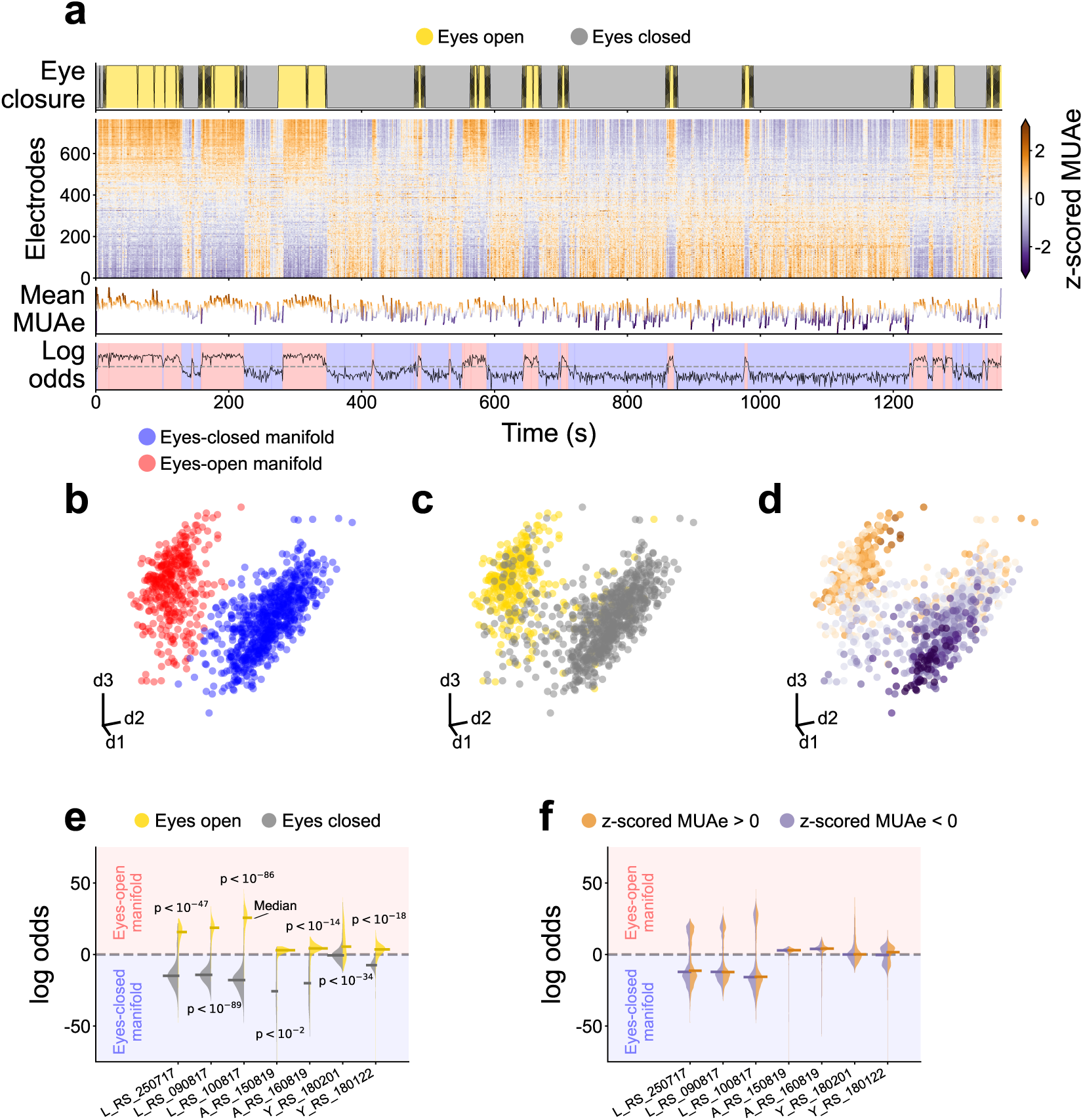
Two distinct neural manifolds in V1 correlated with eye closure. **a** Overview of the experimental data from session L RS 250717. From top to bottom: Time evolution of the eye signal; the z-scored MUAe signal for each electrode (electrodes ordered by their correlation with the eye signal); the mean z-scored MUAe at each time point; and the log odds overlaid with the most likely manifold (two clusters, Gaussian mixture model). **b, c, d** First three principal components of the MUAe population activity. Colors indicate the manifold identified via the log odds of a Gaussian mixture (**b**), the eye closure (**c**) and the mean z-scored MUAe (**d**). Each dot represents a different point in time. Outliers were excluded from the neural manifolds shown in **b**–**d**, see Outlier removal. **e, f** Violin plots of the distribution of the log odds across epochs, respectively distinguished according to eye closure (**e**, result of a logistic regression test shown) and z-scored MUAe (**f**). Horizontal bars indicate medians of the distributions.

In V1, at least two distinct neural manifolds are apparent in the 3D projection space; sample session in Figure 2b-d, see Figure S1-S6 for all other sessions and subjects. We labeled the manifolds according to the sign of the log odds of a two-component Gaussian mixture model (see methods Neural manifolds and clustering and Outlier removal). The log odds represent the probability for a given data point to correspond to one manifold or the other.

To confirm that the two manifolds in the lower-dimensional projection are not an artifact of the dimensionality reduction, we estimated the Betti numbers of the high-dimensional population activity using persistent homology (see Methods, Topological data analysis). The persistence barcodes show that at least two independent generators of the H_0_ homology groups exist in the high-dimensional population activity, corresponding to two connected components (Figure S7), i.e., two distinct neural manifolds. Thus, we confirm that the two manifolds observed in the 3D projection are inherent to the high-dimensional space.

Additionally, we tested whether the observed manifolds could be an artifact of the MUAe signal. We spike-sorted one session (L RS 250717) with a semi-automatic method and analyzed the population activity resulting from the single-neuron firing rates (Figure S8). The spiking activity also displayed two manifolds, in agreement with the MUAe signals.

While the activity of visual cortex is mainly driven by visual input, whether and to what extent it is separately modulated by eye closure is unclear. Marking data points on the V1 manifolds with the eye closure signal (Figure 2b) reveals that one manifold strongly relates to the eyes-open periods, whereas the other manifold strongly relates to the eyes-closed periods.

To confirm the correlation between eye closure and manifolds, we tested the differences between the eyes-open and eyes-closed periods using a twofold approach. First, we performed a logistic regression between the eye closure signal and the log odds, revealing a significantly higher than chance correlation in all sessions (Figure 2e). Second, we visualized the distribution of the log odds during the eyes-open and eyes-closed periods separately; showing a clear correspondence between the eye closure and the sign of the log odds in most cases (Figure 2e). Taken together, the logistic regression and the log-odds distributions demonstrate that membership of a point in state space in one of the two V1 manifolds is closely related to eye closure. Given this, we will refer to the manifolds as the eyes-open manifold or the eyes-closed manifold.

The existence of two separate manifolds could be trivially explained if the MUAe activity levels were significantly higher in one manifold, and the manifolds simply reflected the population activity level. To rule out this possibility, we checked whether higher-activity epochs uniquely correspond to one of the manifolds. The violin plots of the full data distribution—based on the z-scored MUAe shown in Figure 2a—show that there is no clear separation into two manifolds (Figure 2f). Additionally, we visualized the 2D histograms of z-scored MUAe against log odds (Figure S9). Both the violin plots and the 2D histograms suggest that the activity level alone does not fully explain the presence of the two neural manifolds in macaque V1.

For completeness, we also visualized the population activity from V4 and DP (Figure S10, S11). In contrast to V1, the population activity in areas V4 and DP does not appear to contain two distinct neural manifolds. We also tested the relationship between neural activity and eye closure in V4 and DP (Figure S12), using the same procedure as for V1. Although some correlation is observed between eye closure and log odds, the violins reveal no clear manifold separation. Thus, we conclude that the observed manifolds are restricted to V1 and are not present in V4 or DP.

### Higher dimensionality during eyes-open periods, primarily due to decorrelation

To further understand the functional role and implications of the observed neural manifolds in V1, we studied their dimensionality. We used the participation ratio (PR, Equation 1), which is defined via the percentage of variance explained by the principal components of the covariance matrix^19,39^. The PR can be rewritten in terms of the statistics of the covariance matrix

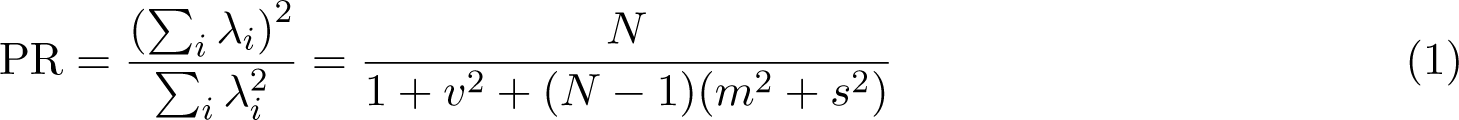

matrix and *N* is the number of electrodes. *v*, *m*, and *s* are the ratios between the standard deviation of auto-covariances, average cross-covariances, and the standard deviation of cross-covariances with respect to the average auto-covariances, respectively. See Dimensionality for detailed methods.

To study the dimensionality, we computed the time-varying PR from the z-scored MUAe signals, by calculating the PR for windows of 30 s width (1 s steps, thus 29 s of overlap with adjacent windows), see Figure 3a. Stronger MUAe activity is typically associated with higher variance, which may bias the results toward higher dimensionality. We avoided bias due to the varying activity level by normalising the data (via z-scoring) within each window. We found that there is a strong correlation between the log odds and the time-varying PR (Figure 3b) and compared the PR values between the two manifolds using a Mann-Whitney U test (Figure 3c). We also measured the PR for one spike sorted session (Figure S8d). The correlation and tests show that the dimensionality is higher during the eyes-open periods, consistently for all datasets, both for MUAe and spikes.

**Figure 3:**
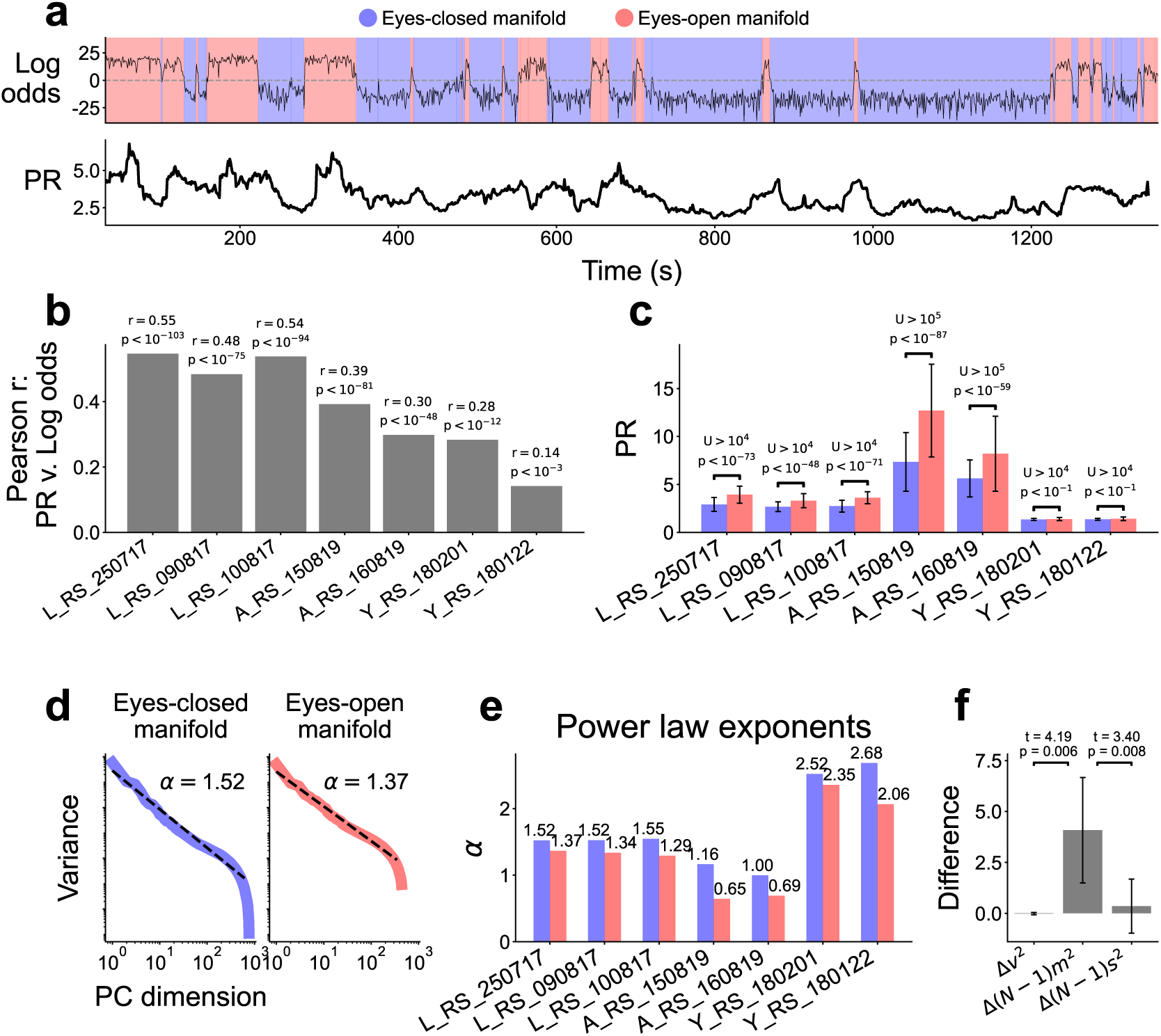
Higher dimensionality during eyes-open periods. **a** Log odds and participation ratio (PR) for session L RS 250717. The PR was calculated on a sliding window of 30 s width. **b** Pearson correlation between log odds and PR for each session. **c** Comparison of PR between neural manifolds (Mann-Whitney U test). **d** Distribution of principal components and their explained variance on a log-log scale, for each manifold. Power law exponent *α* estimated over the ranges where the curves approximate a power law. **e** Comparison of power law exponents for the two neural manifolds in all sessions. The eyes-open manifold always had a smaller exponent, indicating a higher dimensionality. **f** Differences in the terms of the PR function between eyes-open and eyes-closed, results of Welch’s t-test across sessions shown. The quantities are related to the standard deviation of auto-covariances (*v*^2^), average cross-covariances ((*N* − 1)*m*^2^), and standard deviation of cross-covariances ((*N* − 1)*s*^2^)

To further support this finding, we also show the distribution of the variance explained by each of the principal components (PC) of the MUAe data, depicted on a log-log scale in Figure 3d. We fitted a power law to the PC variances and report the exponent *α* (Figure 3e). A higher *α* indicates faster decay of the curve, i.e., lower dimensionality. The power law exponents are in agreement with our sliding window approach: We observe higher dimensionality during eyes-open than during eyes-closed periods (Figure 3e).

To narrow down the reason causing the dimensionality changes, we computed *v*^2^, (*N* − 1)*m*^2^, and (*N* − 1)*s*^2^ and observed that the changes in (*N* − 1)*m*^2^—i.e. the average cross-covariances—dominate the PR differences between the eyes-open and eyes-closed periods (Figure 3f). Thus, the main reason for the observed dimensionality changes is decorrelation during the eyes-open periods.

### Top-down signals from V4 to V1 are present in the form of beta-band spectral Granger causality

In search of an internal mechanism that may modulate the neural manifolds and their dimensionality, we turned our attention to cortico-cortical interactions. Since signatures of top-down activity have previously been reported in the beta frequency band (roughly 12–30 Hz)^31,33,45^, we use spectral Granger causality to measure top-down signals.

To determine whether top-down signals are present in our data, we calculated the coherence and Granger causality between every pair of V1-V4 and V1-DP electrodes (see Coherence and Granger causality)—using the local field potential (LFP). Figure 4a,b show the coherence and Granger causalities for a sample pair of electrodes. To quantify the cortico-cortical signals, we searched for peaks in the coherence and Granger causality, using an automatic method (see Methods, Peak detection).

**Figure 4:**
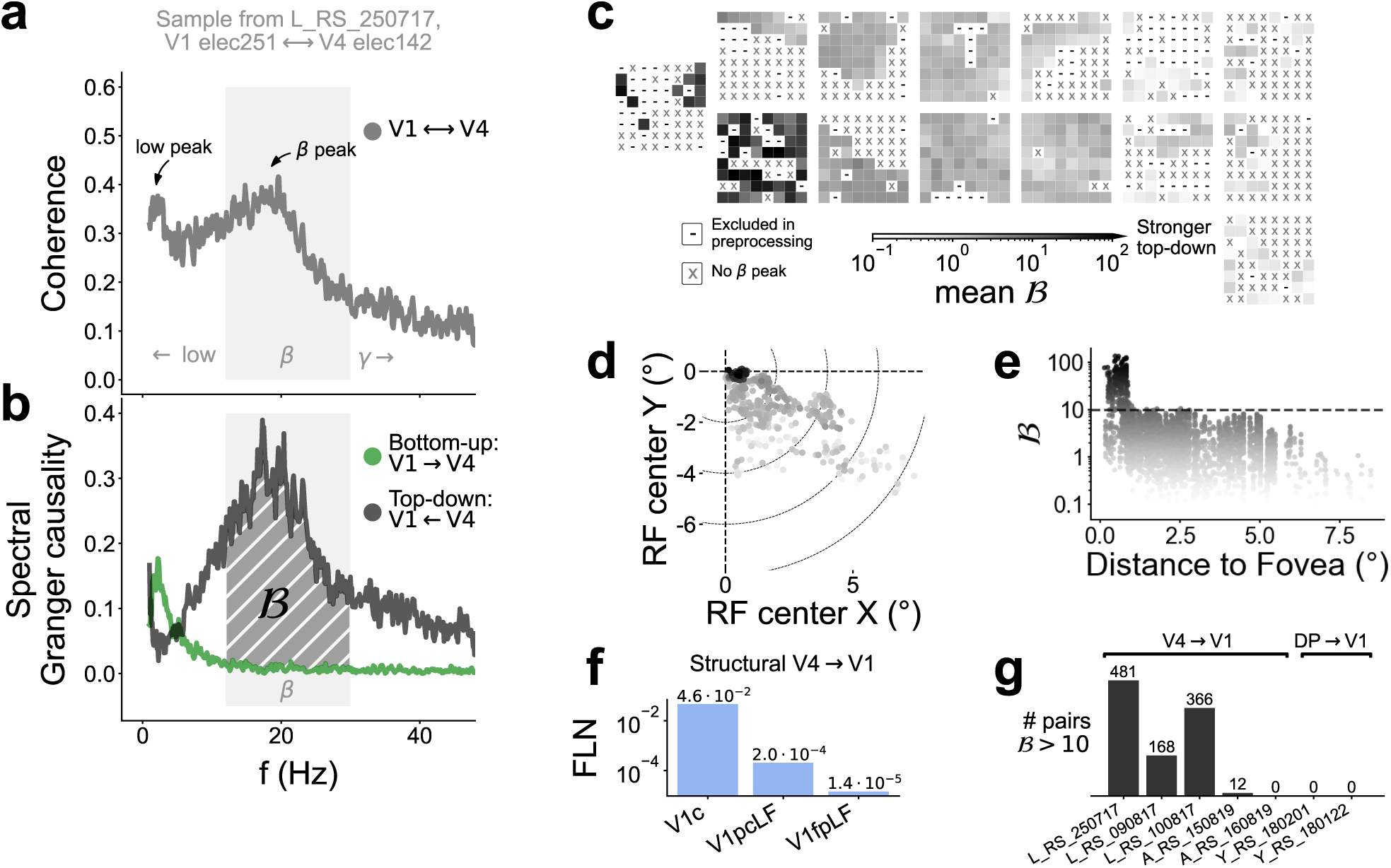
Inter-area coherence and spectral Granger causality. **a** Representative sample of coherence between V1 and V4 (electrodes 242 and 142, respectively). Low-frequency and beta-band peaks indicated. **b** Representative sample of spectral Granger causality. **c** Schematic representation of the electrode locations overlaid with the mean top-down signal strength per electrode (see Coherence and Granger causality for a description of). **d** Receptive field (RF) map overlaid with the mean per electrode. Stronger is found around the foveal region of V1. **e** Mean displayed against the distance from the fovea. **f** Fraction of labeled neurons (FLN) from V4 to V1 (data from tract-tracing experiments ^44^). V1 subdivisions represent c: central (foveal region), LF: lower visual field, pc: peri-central, and fp: far periphery. The strongest connectivity exists from V4 to V1c, in agreement with our measurements. **g** Number of electrode pairs with strong (B *>* 10) top-down signals detected in each session.

We detected beta-frequency Granger causality peaks in around 0.5% of all V1-V4 electrode pairs, predominantly in the top-down direction (Figure S13). We only found beta-band bottom-up interactions in V1-DP electrode pairs.

For the electrodes with a beta causality peak, we estimated the causality strength B (Equation 5). The electrodes with their receptive field (RF) closer to the fovea show substantially higher (Figure 4c–e, Figure S14), in agreement with a previous structural connectivity report ^44^ (Figure 4f). To disregard potential spurious Granger causality peaks, we restrict all further analysis of the top-down signals to the strongest interactions, by setting a threshold of B *>* 10 (Figure 4g). We found few bottom-up V1-to-V4 signals with high strength in the beta frequency band.

We thus found top-down signals from V4 to V1, in agreement with previous studies^31,33,45^; but we did not find strong signals from DP to V1 in our data. V4-to-V1 signals are therefore strong candidates for the modulation of the neural manifolds and their dimensionality.

### Stronger top-down signals from V4 to V1 during eyes-open periods

To elucidate the behavioral relevance of the V4-to-V1 top-down signals, we examined how the LFP spectral power, coherence, and Granger causality change in relation to eye closure.

We extracted the LFP data for each behavioral condition and concatenated the data within the same condition. This approach could introduce some artifacts, which we expect to be minor in view of the very small number of transitions in comparison with the number of data samples (500 Hz resolution). Both in V4 and V1, we find that the spectral power at low frequencies (*<* 12 Hz) is higher during the eyes-closed periods, whereas the power in the beta band (12–30 Hz) is slightly higher during eyes-open periods (Figure 5a, Figure S15). Spectrograms of the V1 LFP power confirm the reduction in low-frequency power during eyes-open periods (Figure S17). The coherence in the beta band is higher during the eyes-open periods, with the peak shifted to higher frequencies compared to the eyes-closed condition. Notably, the top-down Granger causality is substantially higher in the beta band during the eyes-open periods.

**Figure 5:**
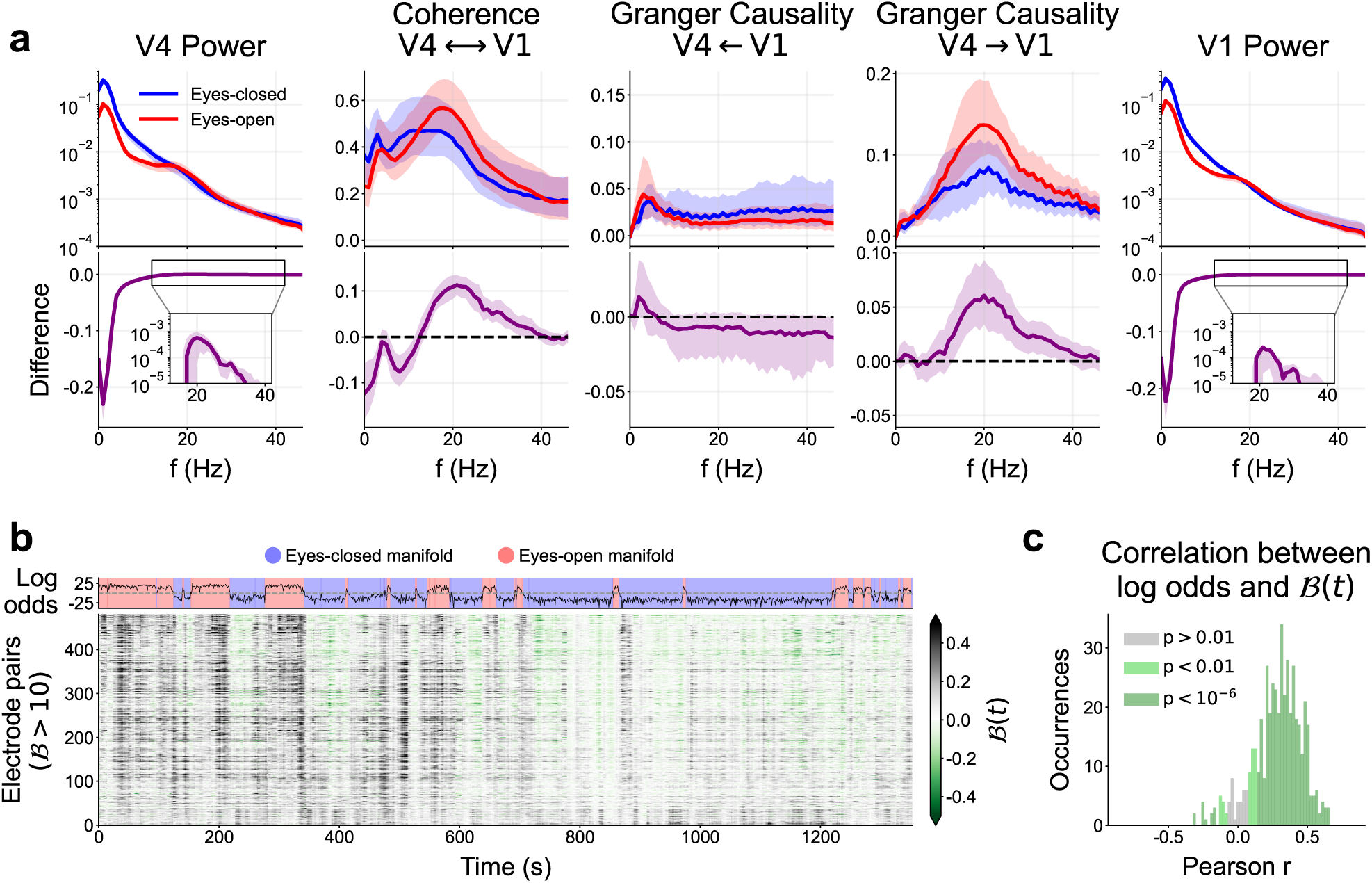
Stronger top-down signals from V4 to V1 during eyes-open periods. **a** Spectral power, coherence, and Granger causality of the LFP for the electrodes with high causality strength ( *>* 10) in session L RS 250717, see Figure S15 for all other sessions. The data for each behavioral condition (eyes-open/closed) were concatenated and their metrics reported separately (top row). The difference between eyes-open and eyes-closed periods was calculated for each electrode or pair of electrodes (bottom row). In all panels the thick line shows the median across electrodes (or pairs of electrodes) and shading indicates the 25th to 75th percentile. **b** Time evolution of log odds (top) and time-dependent beta-band Granger causality difference (t) (bottom), for the electrode pairs with top-down signals. **c** Histogram of the Pearson correlation between the log odds and (t). Color indicates the significance levels of the associated two-sided t-test.

In order to confirm our observations, we also computed spectrograms of the Granger causality using a 10-second sliding window (Figure S16a). Statistical tests (Welch’s t-test) of the difference between bottom-up and top-down Granger causality, ΔGC, confirmed a shift toward top-down interactions during the eyes-open periods compared to the eyes-closed periods, for a vast majority of all electrode pairs (Figure S16b,c). Thus, we found higher beta-band Granger causality during eyes-open periods using two different approaches.

Additionally, to confirm the interdependence of top-down signals and the neural manifolds, we computed the correlation between the time-varying beta-band Granger causality B(t) (Equation 8) and the log odds (Figure 5b,c). An overwhelming majority of V1-V4 electrode pairs showed a highly significant correlation (*p <* 10^−6^, two-sided t-test). Thus, the top-down signals and neural manifolds are co-dependent at a fine temporal scale, as well as within eyes-closed and eyes-open periods.

We further tested whether the top-down signals were correlated with gaze direction and eye movements (Figure S18), to rule out the presence of any visual stimuli—despite the experiments being performed in a dark room. No clear trend could be observed, thus indicating no relation between gaze direction and top-down signals. This finding suggests that the visual scene is not the source of the observed top-down signals.

In conclusion, the time-dependent spectral analysis reveals large variations of power and Granger causality. On the one hand, the spectral power at low frequencies decreases during eyes-open periods, consistent with the well-known alpha blocking phenomenon ^26–28^. On the other hand, the V4-to-V1 top-down signals are strongest during the eyes-open periods. The time-varying top-down beta causality strength did not substantially correlate with gaze direction or eye movements, suggesting no relation between the top-down signals and the visual scene; as expected in a dark room. Taken together, these results suggest that V4-to-V1 signals modulate V1 activity, contributing to a different state-space manifold with increased dimensionality.

## Discussion

In this paper, we presented three novel findings in the primary visual cortex (V1) of macaques during the resting state: two separate manifolds in the state space associated with eye closure (Figure 2); higher dimensionality due to lower mean cross-correlations during eyes-open periods (Figure 3); and the presence of stronger top-down signals from V4 to V1 during the eyes-open periods, primarily targeting the foveal region of V1 (Figure 4, Figure 5). In addition, we observed lower power at frequencies below 12 Hz during the eyes-open periods (Figure 5, Figure S17), consistent with the well-known alpha blocking effect ^26^.

We observed that two distinct manifolds appear in the state space of macaque V1—but not V4 nor DP—during the resting state for all subjects and sessions, for both MUAe and spike data (Figure 2, S1-S6, Figure S8), and are correlated with eye closure (Figure 2e,f). The manifolds were not just an artifact of the three-dimensional projection used for visualisation, as we confirmed they also exist in higher dimensions with persistent homology (Figure S7). Previous work in mice has shown that the visual cortex represents a myriad of behaviors in the resting state, such as facial movements or running^46^. However, a similar study on the macaque showed that the macaque visual cortex is very specific to vision, and minimally driven by spontaneous movements^47^. Thus, we do not expect the neural manifolds of V1 to be strongly affected by any behavior other than visual behavior, in agreement with our finding that eye closure neatly explains the two manifolds.

Our findings could in principle be explained by the presence of complex visual stimuli that would alter the population dynamics and cortico-cortical communication. However, we are certain that no strong visual stimuli are present in the visual field, due to the very dark environment of the recording room. Additionally, we performed several analyses to control for activity levels (Figure 2 and Figure S9) and gaze direction Figure S18. Furthermore, the original data for Macaques L and A includes an extensive evaluation of data quality, which excluded all electrodes that did not strongly respond to visual stimuli^42^. Thus, all the electrodes included in our analysis (from Macaques L and A) would strongly respond if there were strong visual stimuli, but we observed no such responses in the MUAe activity (see Figure 2). We are therefore certain that the visual input is faint or nonexistent, which implies that the observed neural manifolds must be induced by some other internal mechanism.

Further characterisation of the activity in the different manifolds revealed that the neural dimensionality is manifold-dependent (Figure 3). We observed higher dimensionality in the eyes-open manifolds across all macaque and sessions. Our measured dimensionality is in agreement with previous reports on the visual cortex^10,16^. Previous work has also shown higher dimensionality in the primary motor cortex during eyes-open than eyes-closed periods^48^, analogous to our findings in the visual cortex.

We hypothesized that top-down signals from higher cortical areas could be the modulatory mechanism responsible for the changes observed in the neural manifold and dimensionality of V1 activity. Indeed, we found that there are strong top-down signals from V4 to V1 (Figure 4), targeting particularly the foveal region of V1, in agreement with structural connectivity^44^. We also found the top-down signals to vary over time, with increased presence during the eyes-open periods (Figure 5). In agreement with our findings, previous studies found that cortico-cortical top-down signals between V1 and V4 are predominantly present in the beta (12–30 Hz) frequency band, while bottom-up signals between V1 and V4 are present in the delta/theta (*<* 8 Hz) and gamma (*>* 30 Hz) bands^31,45^. Others suggest that top-down signals from V4 to V1 are found more generally in the low frequencies (*<* 30 Hz), not uniquely in the beta-band^33^. In our analysis we did not find any gamma band causality (Figure 4), likely because our recordings were from the deep cortical layers (in macaque L and A the electrodes were 1.5 mm long, putatively recording mostly from layer 5) and gamma oscillations are known to be weak in layer 5 of the visual cortex^30,49^. In addition, gamma activity is associated with bottom-up signals^33^, which we do not expect in a dark room with no visual stimuli. In contrast to our findings, van Kerkoerle et al.^30^ reported that top-down signals appear in the alpha (8–12 Hz) frequency range. Whether the specific top-down and bottom-up frequencies generalize to the whole cortex is unclear. Instead, Vezoli et al. ^45^ postulate overlapping modules of certain frequencies (alpha, low-beta, high-beta, and gamma) that differ across cortical areas. Our findings are also consistent with the work by Semedo et al. ^50^, who suggested that bottom-up signals dominate during visual stimulation and top-down signals dominate in the absence of visual stimuli—note that in their work the eyes were always open.

The spatial organisation of the top-down signals is in agreement with predictions made by the *central-peripheral dichotomy* (CPD) theory^51,52^. In this theory, the central vision is proposed to be primarily concerned with object recognition, and thus should be more strongly targeted by top-down inputs (e.g. from V4). These top-down signals would query additional visual information, ultimately reducing ambiguity in visual processing.

We did not find top-down signals from DP to V1, possibly due to the electrodes used in macaque Y being 1 mm long, thus likely recording from layer 4, and top-down connections do not originate in nor target layer 4 of visual cortex^53^. An alternative explanation might be formulated in the context of the CPD theory, which asserts that top-down inputs in the dorsal stream should preferentially target the far periphery of V1^54^, which our experiments did not record from.

In the present study, it was not possible to test directly from the experimental data whether the V4-to-V1 signals are responsible for the modulation of V1 dynamics. Future studies could perform such a test by a reversible inactivation of the V4-to-V1 pathway, such as via reducing the temperature of V4^55,56^, injecting a GABA agonist (e.g., muscimol, bicuculline)^57–59^ or using targeted optogenetic suppression^60^. These techniques have been successfully applied to study the suppression of cortico-cortical communication; however, to best of our knowledge, they have not been used to study the effects of macaque V4-to-V1 signals in the resting state.

Numerical simulations offer an alternative approach to study the effect of top-down signals in spiking neural networks. We thus performed preliminary simulations of a simple spiking neuron model—of the well-known Brunel type ^61^—to ascertain whether V4-to-V1 signals can modulate the neural manifolds (Figure 6). Modeled top-down signals, in the form of sinusoidal oscillating inhomogeneous Poisson processes, led to a different neural manifold in the network activity when a subset of the network neurons was targeted (Figure 6d). These changes were not due to the increase in firing rate caused by the additional top-down input, but rather due to the activation of different neuron patterns in the model (Figure 6e,f). We limited the analysis of the model to the presence of neural manifolds, because our model was ill-suited to study the dimensionality, given that average cross-correlation is known to cancel out in balanced EI networks^62,63^. Future work could use more complex models—such as clustered networks ^62,64,65^—to study the effects of correlated inputs with realistic power spectra on the dimensionality and elucidate whether the top-down signals can directly induce the observed increase in the dimensionality during the eyes-open periods.

**Figure 6:**
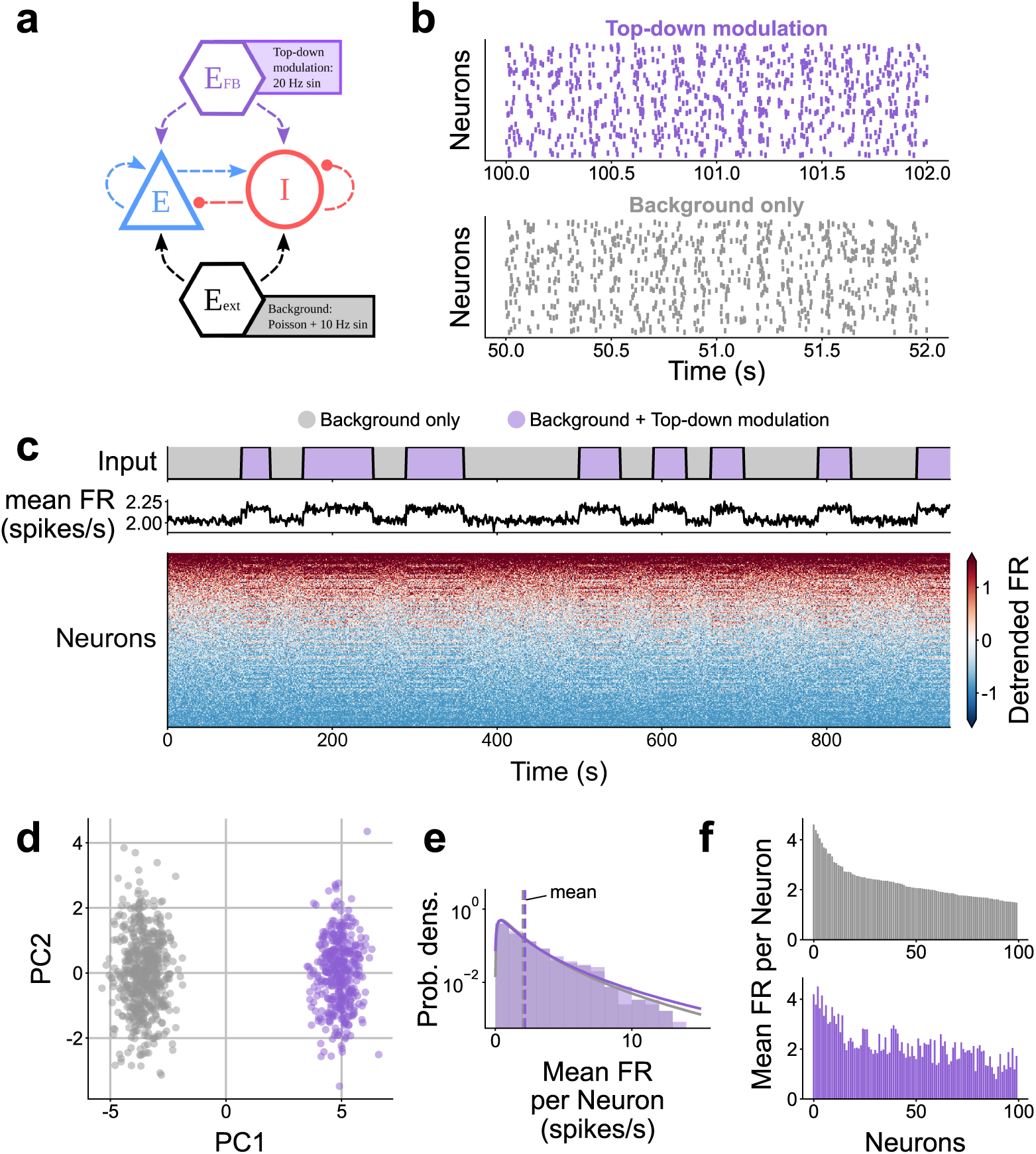
Simulation of a balanced spiking neural network with top-down modulation. **a** Diagram of balanced random spiking neural network. Background input is provided constantly, top-down signals are provided intermittently. **b** Sample raster plots show spiking activity in the different input regimes. **c** Time evolution of input regimes and mean firing rate (FR). **d** First two principal components of the firing rate (binsize = 1 s). Colors indicate the different input regimes. **e** Distribution of mean firing rate per neuron is almost identical between the two regimes. **f** Mean firing rate of the 100 most active neurons. The top-down modulation changes the mean firing rates of each neuron, in both the positive and negative directions, leading to the observed distinct manifolds.

Taken together, our data analysis and simulations suggest that top-down modulation alone is sufficient to cause the distinct neural manifolds in V1 activity. Nevertheless, sustaining the different V1 manifolds might involve additional mechanisms, such as neuromodulation or adaptation of recurrent connectivity via short-term plasticity. Previous work suggests that N-methyl-D-aspartate (NMDA) receptors are central to the top-down communication from V4 to V1^35,66^. Interestingly, targeted pharmacological deactivation of NMDA receptors in macaque V1 leads to the suppression of alpha blocking^28^ and absence of decorrelation during eyes-open ^67^; both of which are correlated with the increased V4-to-V1 signals in our data. In addition, the top-down signals are not constant throughout the eyes-open periods (Figure S16), but the slow timescale of the NMDA receptors could help sustain the eyes-open manifold, even if the top-down input fades. Thus, we speculate that the top-down connections preferentially target the NMDA receptors in V1 neurons, leading to the observed alpha blocking and decorrelation. Another mechanisms could involve recurrent connectivity, in the form of cell-type-specific motifs. Such motifs have been shown to affect the dimensionality of brain networks ^39^, and they could emerge in the effective connectivity of the network as a result of the top-down input.

If the V4-to-V1 signals convey behavioral information, then how does such behavioral information reach V4 in the first place? We explore the possible communication pathways that lead to the observed V4-to-V1 signals, illustrated in Figure 7. We identified three main candidates: the visual stimulus (or absence thereof) from the retina to V1; the proprioception of eyelid muscles via the somatosensory cortex; and the voluntary motor commands for eye closure. The first proposed pathway involves visual stimuli being transmitted from the retina to V1 via the lateral geniculate nucleus (LGN). The absence of stimuli could be the reason for the observed changes in the V1 activity, whereas the presence of visual stimuli could trigger a V1-V4 feedback loop. However, the macaques in our experiments had very little to no visual input, even during eyes-open periods, since they were sitting in a dark room. Additionally, we found no consistent difference in MUAe activity levels between the eyes-open and eyes-closed manifolds (Figure 2). The second proposed pathway involves proprioception of the eyelid that could inform the cortex when the eyes are closed and trigger the activity changes in V1. Mechanoreceptors in the eyelid activate the oculomotor nerve projecting to the midbrain (possibly to the superior colliculus)^68^, eventually entering the cortex via the somatosensory area (S1)^69^. From S1 the signal could find its way to V1 via several cortico-cortical pathways, potentially including neurons in V4; however this mechanism might be relatively slow, given the absence of direct connections from S1 to V1 or V4^70^. Furthermore, the shortest known S1-to-V1 cortico-cortical pathway does not involve V4, rendering this pathway a rather weak candidate. The third and final proposed pathway involves voluntary eyelid closure which is initiated by the ventral motor cortex and the frontal eye field (FEF). The eyelid closure and eye movements may be communicated to the visual cortex via cortico-cortical connections or the superior colliculus. Given that V4 is part of the fronto-parietal network (with strong FEF⇄V4 connections) ^70,71^, the eye movement signals could easily reach V4, which could then modulate the V1 activity. A trans-thalamic pathway through the pulvinar could also assist the V4-to-V1 communication, including the synchronisation of the alpha rhythm^72^; although such trans-thalamic connections have only been confirmed in mice so far^73^.

**Figure 7:**
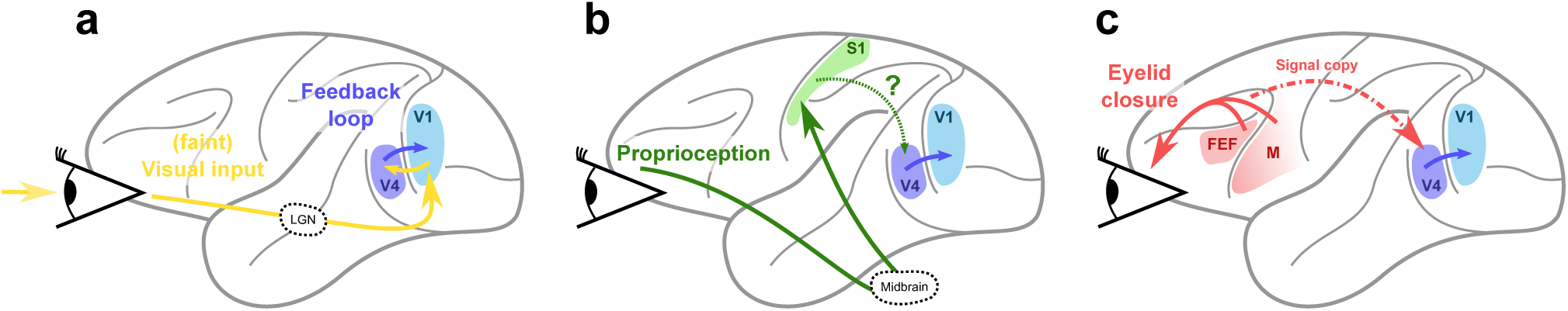
Proposed communication pathways for V1 modulation via V4. **a** Visual input directly to V1 triggering a cortico-cortical feedback loop. **b** Proprioception of eyelid muscles via the midbrain (possibly superior colliculus) and somatosensory cortex. **c** Cortico-cortical communication of motor commands.

The hypotheses from Figure 7 are not necessarily mutually exclusive, and could all play a role in the modulation of V1 activity. To understand which pathways are most relevant to sustain the manifolds, we had a closer look around the manifold transitions (Figure S19) by looking at the MUAe signals at a high temporal resolution (1 kHz). For the eye-opening transitions, we observed that sometimes V1 MUAe activity precedes V4 activity, in agreement with the feedback loop hypothesis (Figure 7a); whereas in other cases V4 activity precedes V1 activity, in agreement with the hypotheses shown in Figure 7b,c. In the eye-closing transitions, the activity from V1 and V4 appeared to be simultaneous. The number of transitions was relatively small, which did not allow for a quantitative analysis of the transitions between the two manifolds. Further work could revisit this issue by looking at longer recordings including larger numbers of transitions between eyes-open and eyes-closed periods.

Given the complex mechanisms that seem to be involved in ensuring that V1 population activity adjusts to eye closure, it seems likely that it has a functional benefit. First of all, if the eyes are closed, no visual stimuli are processed and V1 firing rates are reduced to save energy. On the other hand, when the eyes are open, higher-dimensional activity might be advantageous for better encoding visual stimuli, which are known to have a high dimensionality^10^. This could thus facilitate visual processing. Previous work showed that spectral power in the alpha band (8–12 Hz) is inversely correlated with visual recognition performance in human subjects^27,74^: lower alpha power was associated with better performance in a visual discrimination task. Our results suggest that the change in neural manifolds and dimensionality are directly correlated with the decrease in alpha power (Figure 5, Figure S17). Future work could study the relation between the dimensionality, alpha power, and visual performance (e.g., response latency to different images) to determine the functional relevance of our findings.

In conclusion, we provide *in vivo* evidence for the modulation of neural manifolds by cortico-cortical communication, which we hypothesize could enable more efficient responses to visual stimuli. Our analysis and previous results suggest that the eyes-open manifold—together with the corresponding dimensionality and spectral power changes—constitutes a visual stand-by state, which is modulated by top-down input from V4 and other internal mechanisms.

## STAR Methods

### RESOURCE AVAILABILITY

#### Lead contact

Further information and requests for resources should be directed to and will be fulfilled by the lead contact, Aitor Morales-Gregorio

#### Materials availability

This study did not generate new unique reagents.

#### Data availability

The electrophysiology data from macaques L and A (raw, MUAe, LFP, behavioral) were previously published and are publicly accessible ^42^. The electrophysiology data for macaque Y, together with all processed data for this project have been deposited in a GIN repository and are publicly available at the time of publication under a Creative Commons Attribution-ShareAlike 4.0 International Public License. The DOI to the repository is listed in the key resources table.

All original code has been deposited at the same GIN repository and is publicly available as of the date of publication. The DOI to the repository is listed in the key resources table.

Any additional information required to reanalyze the data reported in this paper is available from the lead contact upon request.

### EXPERIMENTAL MODEL AND STUDY PARTICIPANT DETAILS

#### Macaques

We analyzed the resting state data from three (N=3) rhesus macaques (*Macaca mulatta*), recorded in two different experimental laboratories. The data from macaques L & A was collected at the Netherlands Institute for Neuroscience, and previously published ^42^. The data from macaque Y was collected at the Institut de Neurosciences de la Timone, with the recording apparatus described elsewhere ^43^. At the time of visual cortex array implantation macaque L (male) was 7 years old and weighed 11 kg; macaque A (male) was 7 years old and weighed 12.6 kg; and macaque Y (female) was 6 years old and weighed 7 kg.

All experimental and surgical procedures for Macaque L & A complied with the NIH Guide for Care and Use of Laboratory Animals, and were approved by the institutional animal care and use committee of the Royal Netherlands Academy of Arts and Sciences (approval number AVD-8010020171046).

All experimental and surgical procedures for Macaque Y were approved by the local ethical committee (C2EA 71; authorization Apafis#13894-2018030217116218v4) and conformed to the European and French government regulations.

#### Electrophysiological data from macaques L & A

We used publicly available ^42^ neural activity recorded from the neocortex of rhesus macaques (N=2) during rest and a visual task. The macaques were implanted with 16 Utah arrays (Blackrock microsystems), two of them in visual area V4 and the rest in the primary visual cortex (V1), with a total of 1024 electrodes. The electrodes were 1.5 mm long, thus recording from the deeper layers, likely layer 5. The system recorded the electric potential at each electrode with a sampling rate of 30 kHz. A full description of the experimental setup and the data collection and preprocessing has already been published ^42^; here we only provide the details relevant to this study.

Three resting-state (RS) sessions were recorded per macaque, during which the subjects did not have to perform any particular task and sat in a quiet dark room. Pupil position and diameter data were collected using an infrared camera in order to determine the direction of gaze and eye closure of the macaques. On the same days as the RS recordings, a visual response task was also performed. The visual response data were used to calculate the signal-to-noise ratio (SNR) of each electrode, and all electrodes with an SNR lower than 2 were excluded from further analysis. Additionally, we excluded up to 100 electrodes that contributed to high-frequency cross-talk in each session, as reported in the original data publication^42^. The sessions, duration and number of electrodes per subject are listed in Table 1.

#### Electrophysiological data from macaque Y

In addition to the published data from macaques L & A, we also used an unpublished dataset from one additional rhesus macaque (N=1). Neural activity was recorded during rest and during a visuomotor integration task. The recording apparatus is described elsewhere^43^. Macaque Y was implanted with five Utah arrays (Blackrock microsystems), two of them in the primary visual cortex (V1), one in dorsal prelunate cortex (area DP), one in area 7A and one in the motor cortex (M1/PMd). In this study we only included the 6×6 electrode arrays from V1 (two arrays) and DP (one array), for a total of 108 electrodes. The electrodes were 1 mm long, thus recording from the central layers, likely layer 4. The recording system recorded the electric potential at each electrode with a sampling rate of 30 kHz.

Two resting-state (RS) sessions were recorded, during which the macaque did not have to perform any particular task and sat in a quiet dark room. Pupil position and diameter data were collected using an infrared camera in order to determine the gaze direction and eye closure of the macaque. We excluded up to 50% of the electrodes that contributed to high-frequency cross-talk in each session, similarly to the methods described in^42^. See Table 1 for an overview of the sessions used in this study.

#### MUAe and LFP signals

The raw neural data were processed into the multi-unit activity envelope (MUAe) signal and local field potential (LFP). To obtain MUAe data, the raw data were high-pass filtered at 500 Hz, rectified, low-pass filtered at 200 Hz, and downsampled to 1 kHz. Finally, the 50, 100, and 150 Hz components were removed with a band-stop filter in order to remove the European electric grid noise and its main harmonics. To obtain the LFP data, the raw data was low-pass filtered at 250 Hz, downsampled to 500 Hz and a band-stop filter was applied to remove the European electric grid noise (50, 100, and 150 Hz).

The MUAe and LFP data for macaques L & A were already provided by the original authors ^42^ in the open-source .nix format, which uses python-neo data structures to hierarchically organize and annotate electrophysiological data and metadata. The metadata, such as the cross-talk removal or the positions of the arrays in the cortex, were provided in the .odml machine-and human-readable format, which were incorporated into the python analysis scripts.

#### Spike sorting

The raw data from one session (L RS 250717) were spike-sorted using a semi-automatic workflow with Spyking Circus—a free, open-source, spike-sorting software written entirely in Python^75^. An extensive description of the methods of this algorithm can be found in their publication, as well as in the online documentation of Spyking Circus^1^.

Roughly, Spyking Circus first applied a band-pass filter to the raw signals between 250 Hz and 5 kHz. Next, the median signal across all 128 channels that shared the same reference (2 Utah arrays) was calculated and subtracted, in order to reduce cross-talk and movement artifacts. The spike threshold was set conservatively, at eight times the standard deviation of each signal. After filtering and thresholding, the resulting multi-unit spike trains were whitened—removing the covariance from periods without spikes to reduce noise and spurious spatio-temporal correlations. After whitening, a subsample of all spike waveforms is selected, reduced to the first five principal components, and clustered into different groups with the k-medians method. Finally, all spikes in each electrode are assigned to one of the waveform clusters based on a template fitting algorithm, which can also resolve overlapping waveforms. After the automatic sorting, the waveform clusters were manually merged and labeled as single-unit activity, multi-unit activity, or noise. Only single-unit activity (SUA) spike trains were included in this study. The waveform signal-to-noise ratio (wfSNR) was calculated for all SUA, and those with a wfSNR *<* 2 or electrode SNR *<* 2 (from the visual response task) were excluded from the analysis.

### QUANTIFICATION AND STATISTICAL ANALYSIS

#### Neural manifolds and clustering

The MUAe data were downsampled to 1 Hz and arranged into a single array, with between 50 and 900 recording locations per session.

In order to visualize the data, we used a standard dimensionality reduction technique (principal component analysis, PCA) to reduce the neural manifold to 3D. The clusters observed in the RS sessions were labeled using a two-component Gaussian mixture model on the 3D projection. The clustering method provides the log odds, i.e., the chance that any given point belongs to one cluster or the other. The log odds captures the multi-cluster structure of the manifold in a single time series; thus, we consider it to be an identifier of the V1 manifolds.

#### Outlier removal

The neural manifolds in our analysis are a collection of time points scattered across the state space. In the data some time points appear very distant from all other points, which we associate with noise and we therefore seek to remove them. To identify the outliers we used a procedure similar to the one used by Chaudhuri et al. ^5^. First, we calculated the distance matrix of all points to each other, and took the 1st percentile value from the distance distribution, *D*_1_. We then estimated the number of neighbours that each point had within *D*_1_ distance, and finally discarded the 20 percent of points with the fewest neighbours.

#### Topological data analysis

We used persistent homology to confirm that the lower-dimensional structures that we observed in the 3D projection of the neural manifolds are in fact topological features of the data and not just an artifact of the dimensionality reduction. Before computing the persistence barcodes we projected the data into a 10D subspace using the isomap technique^76^. The method aims at approximately preserving the geodesic distance between data points (that is the shortest path between two points on the neural manifold) and thus is suited for reducing the dimensionality of the data when applying a topological data analysis. The analysis on the 10D data showed qualitatively equivalent results to the full-dimensional data, while requiring a much shorter computation time.

To calculate the persistence barcodes of the Vietoris-Rips complex of the neural manifold we used an efficient open-source implementation (Ripser^2^). Briefly, the algorithm successively inflates balls with radius *r* around each point of the manifold. If *k* points have a pairwise distance smaller than *r* (that is, for all pairs of points both points are contained in the ball of the other point), they form a (*k* − 1)D simplex. Thus, the neural manifold gives rise to a simplicial complex (a collection of simplices of potentially different dimension) the topological features of which represent the topology of the neural manifold and can be extracted computationally. As *r* is increased, many short-lived features appear by chance. If the manifold has complex topological structures, they should continuously appear as the radius of the balls grows for a large range of *r*. We computed the persistence barcode for the first three homology groups *H*_0_, *H*_1_ and *H*_2_. Homology groups are topological invariants that capture topological features of a given dimension of the neural manifold. The long-lasting bars in the *n*-th persistence barcode correspond to the number of independent generators *β_n_*of the respective homology group *H_n_*. For low dimensionalities, they can be interpreted intuitively: *β*_0_ is the number of connected components, *β*_1_ the number of 1D holes, *β*_2_ the number of enclosed 2D voids. Throughout all plots of this paper we display the top 1% longest-lasting barcodes for each homology group.

#### Dimensionality

We used two different approaches to study the dimensionality of the neural data.

First, we compute the time-varying participation ratio (PR, Equation 1) from the covariance matrix. We take a 30 s sliding window with a 1 s offset over the MUAe data and compute the PR for each window separately. Higher activity leads to higher variance; thus, we normalized the data within each window via z-scoring to minimize this effect. The PR does not require setting an arbitrary threshold. From the time-varying PR we measured the correlation between the log odds and the PR, and the PR distribution in each manifold.

Second, we computed the eigenvalue distribution of the neural data within each manifold. Once again we normalized the data after sampling each manifold. The distribution appeared to follow a power law, in agreement with previous studies ^10^. We used a linear regression in log-log space to fit a power law to our data, where the slope of the linear fit in the log-log plot corresponds to the exponent *α* of the power law.

#### Coherence and Granger causality

To estimate the communication between cortical areas we rely on the coherence and Granger causality of the LFP.

Coherence is the quantification of linear correlations in the frequency domain. Such that

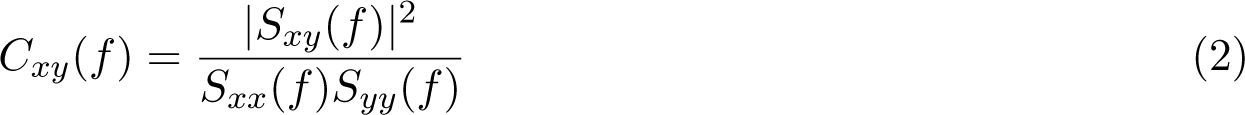

where *C_xy_* is the frequency (*f*) dependent coherence between two signals *x* and *y*, *S_xy_*(*f*) is the cross-spectral density, and *S_xx_*(*f*) and *S_yy_*(*f*) are the auto-spectral densities.

In order to assess the directionality of frequency dependent interactions between the areas we applied spectral Granger causality analysis to the LFP recordings^77^. We first computed the cross-spectral matrix *S*(*f*) with the multitaper method. To this end, we subdivided the chosen signal pairs into 10 s long segments. These were processed individually with 3 Slepian tapers and averaged in the end. This yielded the cross-spectrum. The segments had an overlap of 50%. Next, we decomposed the cross-spectrum into the covariance matrix Σ and the transfer function *H*(*f*) with the Wilson spectral matrix factorisation^78^, obtaining the matrix equation

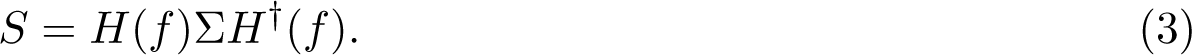

With these factors, one is able to obtain a version of directional functional connectivity between the first and second signals via

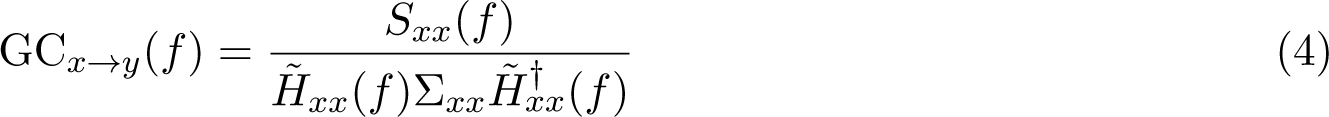

where 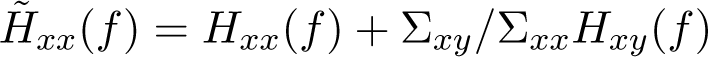 and mutatis mutandis for the influence of the second onto the first signal. The analysis was performed for all pairs of channels between the areas that exhibited a peak in the coherence in the *β* band 12 Hz *< f <* 30 Hz.

We quantify the beta-band Granger causality strength as

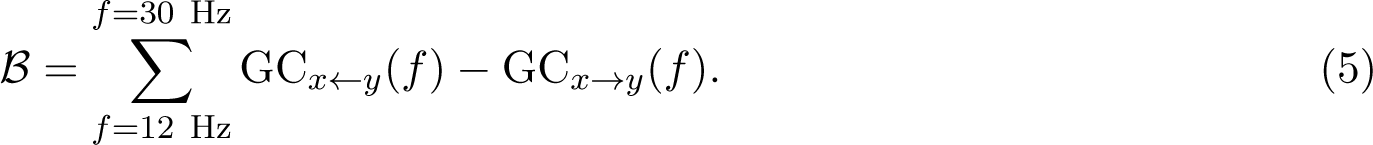

We also analyzed the time-varying spectral Granger causality. For this aim we used 10 s windows and moved them across the data with 1 s steps, for a final time resolution of 1 Hz. We calculated the spectral Granger causality for each window separately. The initial and final 5 s were discarded to avoid disruptions at the boundaries. So the time-varying causality spectrogram is

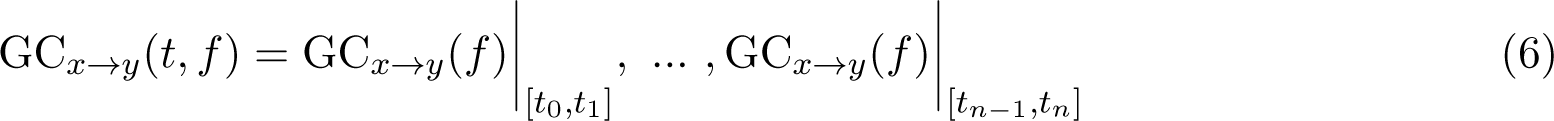

and mutatis mutandis for the *y* → *x* direction.

Finally, we also define the time-varying Granger causality difference

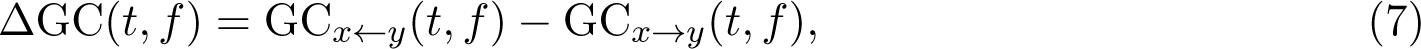

which summed over the beta band we call

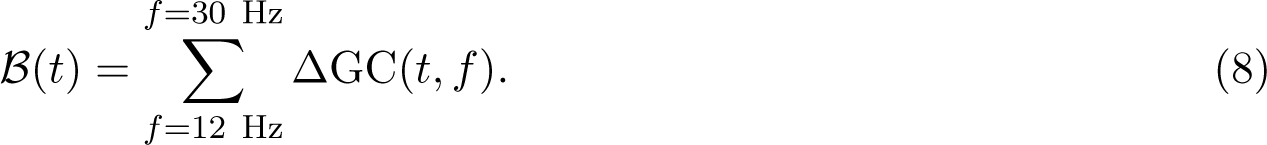

Note that

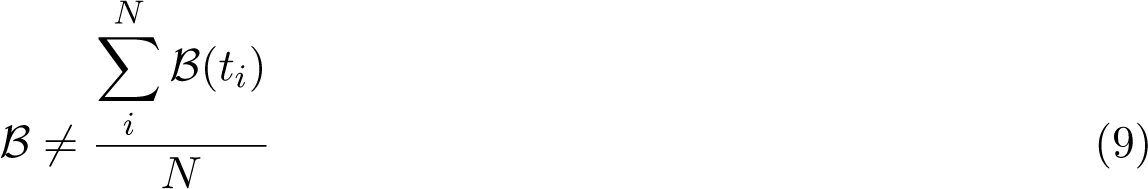

due to the nonlinearities in the Granger causality calculation.

Both the coherence and spectral Granger causality were implemented in the Electrophysiology Analysis Toolkit (Elephant)^79^.

#### Peak detection

To detect coherence and Granger causality peaks (used to identify admissible channel pairs for our analysis) we used a standard peak detection algorithm for 1D arrays using wavelet transforms. We computed the continuous wavelet transform (cwt) for wavelets with widths from 10 Hz to 100 Hz (at 0.1 Hz steps), using a Ricker wavelet—i.e., a Mexican hat. Next, we searched for ridge lines in the cwt—peaks across different wavelet lengths—following standard criteria^80^. Finally, the ridge lines were filtered based on their total length, gaps, and signal-to-noise ratio (snr). The resulting ridge lines (if any) were considered as peaks in the coherence.

The detected peaks tended to be broad, since our parameter choice intentionally rejected narrow peaks. We chose this configuration in favor of robustness and to minimize false positives. Nevertheless, peaks were detected for a majority of electrode pairs.

#### Spiking neural network simulations

To investigate the hypothesis that top-down signals in the *β*-band induce a change in the population dynamics and dimensionality, we conducted a spiking neural network simulation. The network consists of 10, 000 excitatory and 2, 000 inhibitory leaky integrate-and-fire (LIF) neurons with exponential post-synaptic currents. Pairs of neurons are randomly connected with a connection probability of *p* = 0.1. The spike transmission delay is randomly sampled following a log-normal distribution. Generally speaking, the simulation experiments consist of two parts corresponding to the two states observed in the neuronal activity. In the first state (background state), the input consists of spike trains sampled from an inhomogeneous Poisson process with a baseline rate of *ν*_bg_ Hz that is modulated with a 1 Hz sinusoidal oscillation. In the second state, the network additionally receives input spike trains from inhomogeneous Poisson processes with rates oscillating at 20 Hz. The first state represents the eyes-closed, the second the eyes-open condition. Both input regimes provide independent input to each neuron, based on the same rate profiles. During the simulation, we recorded the spiking activity of 1, 000 excitatory and 200 inhibitory neurons. We provided the top-down modulation to a subset of the neurons in the network. We targeted 50% of both the excitatory and inhibitory population. During the simulation, we distinguish two manifolds corresponding to the eyes-open and eyes-closed periods during the recordings. See Table 3, Table 4 for a full description of the network and the experiments. For the simulations we used NEST (version 3.3) ^81^.

**Table 3:** General model description

| **Model Summary** |  |
| --- | --- |
| Populations | two populations, one excitatory, one inhibitory |
| Connectivity | random connectivity |
| Neuron model | leaky integrate-and-fire model |
| Synapse model | alpha-function shaped postsynaptic current |
| Input | independent spike trains from inhomogeneous Poisson processes with given rate $r(t)$ |
| **Neuron and synapse model** |  |
| Subthreshold dynamics | $\frac{dV}{dt} = -\frac{V}{\tau_m} + \frac{I_{\text{syn}}(t)}{C_m},$ $I_{\text{syn}}(t) = J \frac{e}{\tau_{\text{syn}}} (t - t^* - d) e^{-(t-t^*-d)/\tau_{\text{syn}}} \times$ $H(t - t^* - d)$ |
| Spiking | If $V(t-) < \theta$ and $V(t+) \geq \theta$,<br>1. Set $t^* = t$ and $V(t) = V_0$, and<br>2. Emit spike with time stamp $t^*$. |
| **Connectivity** |  |
| Type | pairwise Bernoulli,<br>i.e., for each pair of neurons generate a synapse with probability $p$ |
| Weights | fixed source- and target-population-specific weights |
| Delays | log-normally distributed delays for excitatory and inhibitory neurons |
| **Input** |  |
| Background | $r(t) = \max(0, \nu_{\text{base}}^{\text{bg}} + \nu_{\text{amp}}^{\text{bg}} \cdot \sin(2\pi f^{\text{bg}} \cdot t))$ |
| Top-down modulation | $r(t) = \max(0, \nu_{\text{base}}^{\text{td}} + \nu_{\text{amp}}^{\text{td}} \cdot \sin(2\pi f^{\text{td}} \cdot t))$ |

**Table 4:** Simulation parameters.

| **Population Parameters** |  |  |
| --- | --- | --- |
| $N_{\text{ex}}$ | 10,000 | number of excitatory neurons |
| $N_{\text{in}}$ | 2,000 | number of inhibitory neurons |
| **Connectivity Parameters** |  |  |
| $p$ | 0.1 | connection probability |
| **Neuron parameters** |  |  |
| $\tau_{\text{m}}$ | 20 ms | membrane time constant |
| $\tau_{\text{r}}$ | 2 ms | absolute refractory period |
| $\tau_{\text{syn}}$ | 2 ms | postsynaptic current time constant |
| $C_{\text{m}}$ | 250 pF | membrane capacity |
| $V_{\text{m}}$ | 0 mV | resting potential |
| $E_{\text{L}}$ | 0 mV | membrane capacity |
| $V_{\text{reset}}$ | 0 mV | reset membrane potential |
| $V_{\text{th}}$ | 20 mV | threshold |
| **Stimulus parameters: Background** |  |  |
| $\nu_{\text{base}}^{\text{bg}}$ | 8682 spikes/s | baseline rate |
| $\nu_{\text{amp}}^{\text{bg}}$ | 2170 spikes/s | amplitude |
| $f$ | 10 Hz | sinusoidal oscillation frequency |
| **Stimulus parameters: Top-down signal** |  |  |
| $\nu_{\text{base}}^{\text{td}}$ | 0 spikes/s | base line rate |
| $\nu_{\text{amp}}^{\text{td}}$ | 723 spikes/s | amplitude |
| $f$ | 20 Hz | sinusoidal oscillation frequency |
| $p^{\text{td}}$ | 0.5 | fraction of neurons targeted by top-down modulation |
| **Synapse parameters** |  |  |
| $J_{\text{EE}}$ | 6.4 pA | synaptic efficacy excitatory to excitatory |
| $J_{\text{IE}}$ | 9.5 pA | synaptic efficacy excitatory to inhibitory |
| $g$ | 4 | relative inhibitory synaptic efficacy |
| $J_{\text{EI}}$ | $-g * J_{\text{EE}}$ | synaptic efficacy inhibitory to excitatory |
| $J_{\text{II}}$ | $-g * J_{\text{EE}}$ | synaptic efficacy inhibitory to inhibitory |
| **Delay parameters** |  |  |
| $\mu_{\text{ex}}$ | 1.5 ms | mean of lognormal distribution<br>for excitatory connections |
| $\mu_{\text{in}}$ | 0.75 ms | mean of lognormal distribution<br>for inhibitory connections |
| $\sigma_{\text{ex,in}}$ | 0.5 ms | standard deviation of lognormal<br>distribution for all connections |

### KEY RESOURCES TABLE

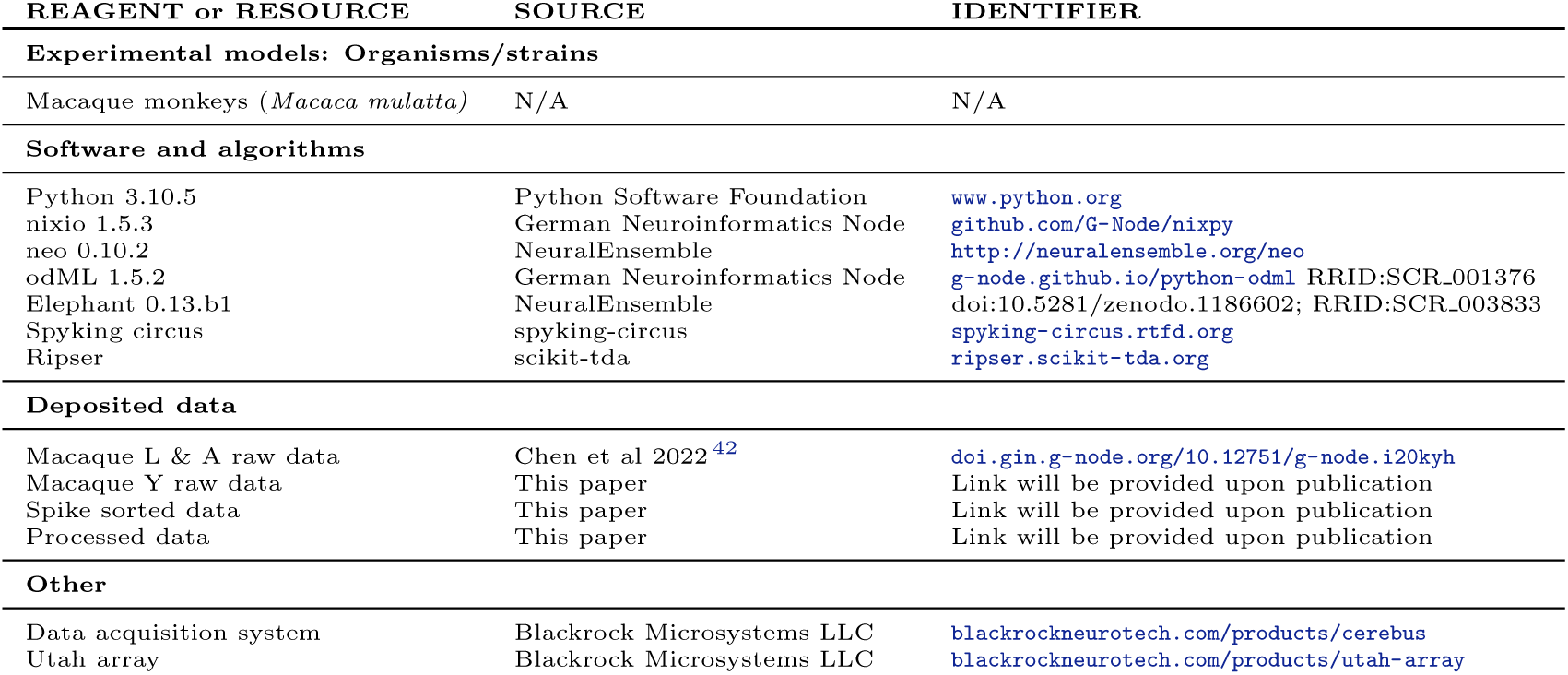

| REAGENT or RESOURCE | SOURCE | IDENTIFIER |
| --- | --- | --- |
| **Experimental models: Organisms/strains** |  |  |
| Macaque monkeys (*Macaca mulatta*) | N/A | N/A |
| **Software and algorithms** |  |  |
| Python 3.10.5<br>nixio 1.5.3<br>neo 0.10.2<br>odML 1.5.2<br>Elephant 0.13.b1<br>Spyking circus<br>Ripser | Python Software Foundation<br>German Neuroinformatics Node<br>NeuralEnsemble<br>German Neuroinformatics Node<br>NeuralEnsemble<br>spyking-circus<br>scikit-tda | www.python.org<br>github.com/G-Node/nixpy<br>http://neuralensemble.org/neo<br>[g-node.github.io/python-odml](https://github.com/g-node/python-odml) RRID:SCR_001376<br>doi:10.5281/zenodo.1186602; RRID:SCR_003833<br>spyking-circus.rtfd.org<br>ripser.scikit-tda.org |
| **Deposited data** |  |  |
| Macaque L & A raw data<br>Macaque Y raw data<br>Spike sorted data<br>Processed data | Chen et al 2022<sup>42</sup><br>This paper<br>This paper<br>This paper | doi.gin.g-node.org/10.12751/g-node.i20kyh<br>Link will be provided upon publication<br>Link will be provided upon publication<br>Link will be provided upon publication |
| **Other** |  |  |
| Data acquisition system<br>Utah array | Blackrock Microsystems LLC<br>Blackrock Microsystems LLC | blackrockneurotech.com/products/cerebus<br>blackrockneurotech.com/products/utah-array |

## Author contributions

AMG, JI, and AK conceptualized the study. AMG curated and processed the data from macaques L and A. AKJ curated and processed the data from macaque Y, with minor inputs from AMG. AMG and AK performed the dimensionality analysis, statistical testing, and Granger causality analysis. AK performed the spiking neuron network simulations and AMG analyzed the results. AMG created all figures, with feedback from all other authors. AMG wrote the initial manuscript; all other authors reviewed the manuscript and provided feedback. JI, SG, and SvA supervised the project, guiding the scope via active discussions. SG and SvA procured the funding and provided the necessary resources.

## Acknowledgements

We thank David Dahmen for his support with the dimensionality analysis and modeling. We thank Simon Essink, Peter Bouss and Tobias Kühn for useful feedback about the manifolds and dimensionality. This project received funding from the DFG Priority Program (SPP 2041 ”Computational Connectomics”) [S.J.van Albada: AL 2041/1-1]; the EU’s Horizon 2020 Framework Grant Agreement No. 945539 (Human Brain Project SGA3); the FLAG-ERA grant PrimCorNet; the DFG (RTG 2416 ”MultiSenses-MultiScales”); the NRW-network ’iBehave’ (grant number NW21-049), and the CNRS

Multidisciplinary Exploratory Projects initiative.

## Declaration of Interests

The authors declare that the research was conducted in the absence of any commercial or financial relationships that could be construed as a potential conflict of interest.

## Extended data figures

**Figure S1:**
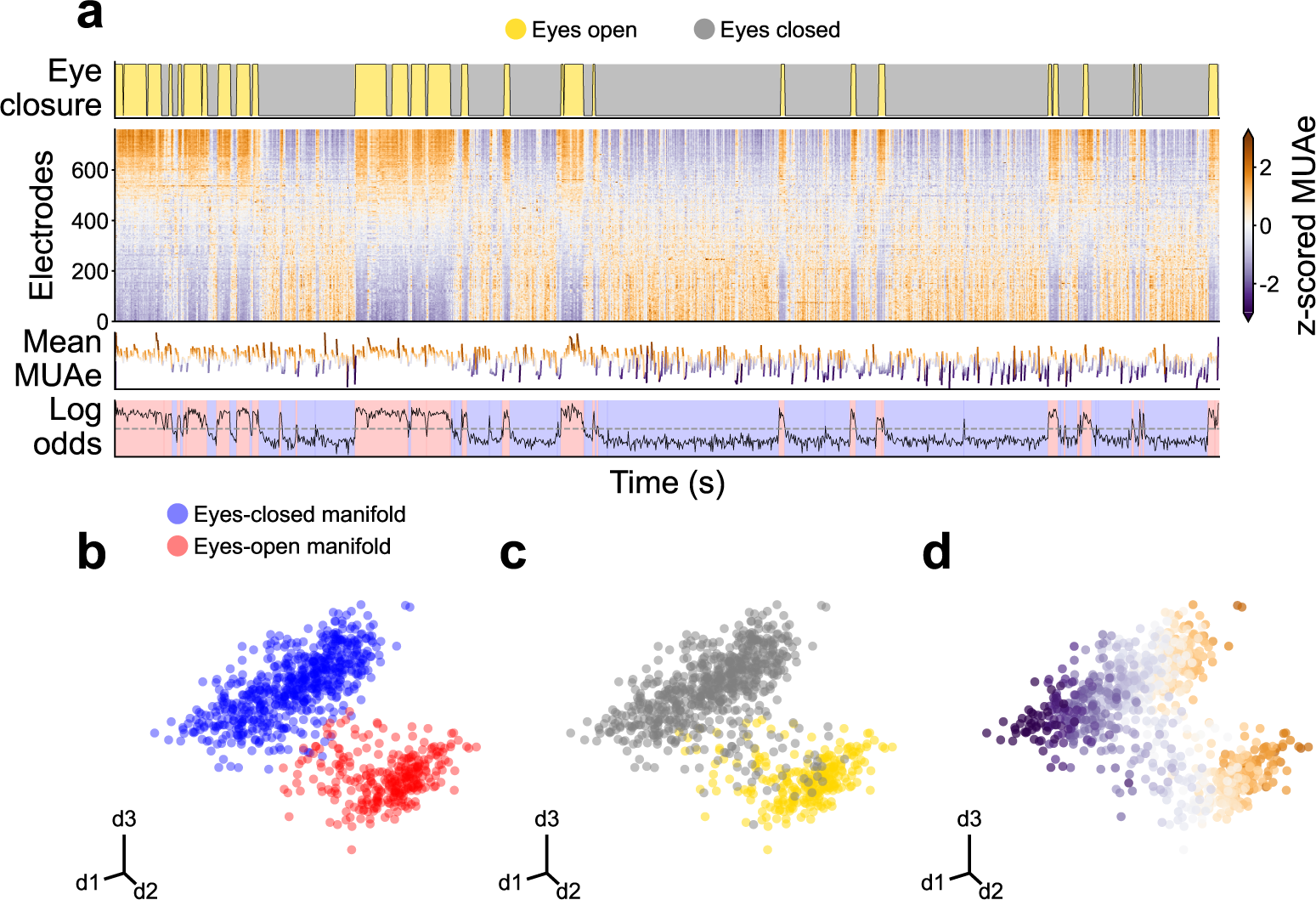
Overview of the experimental data from session L RS 090817. **a** Time evolution of signals. **b, c, d** First three principal components of the MUAe neural manifold.

**Figure S2:**
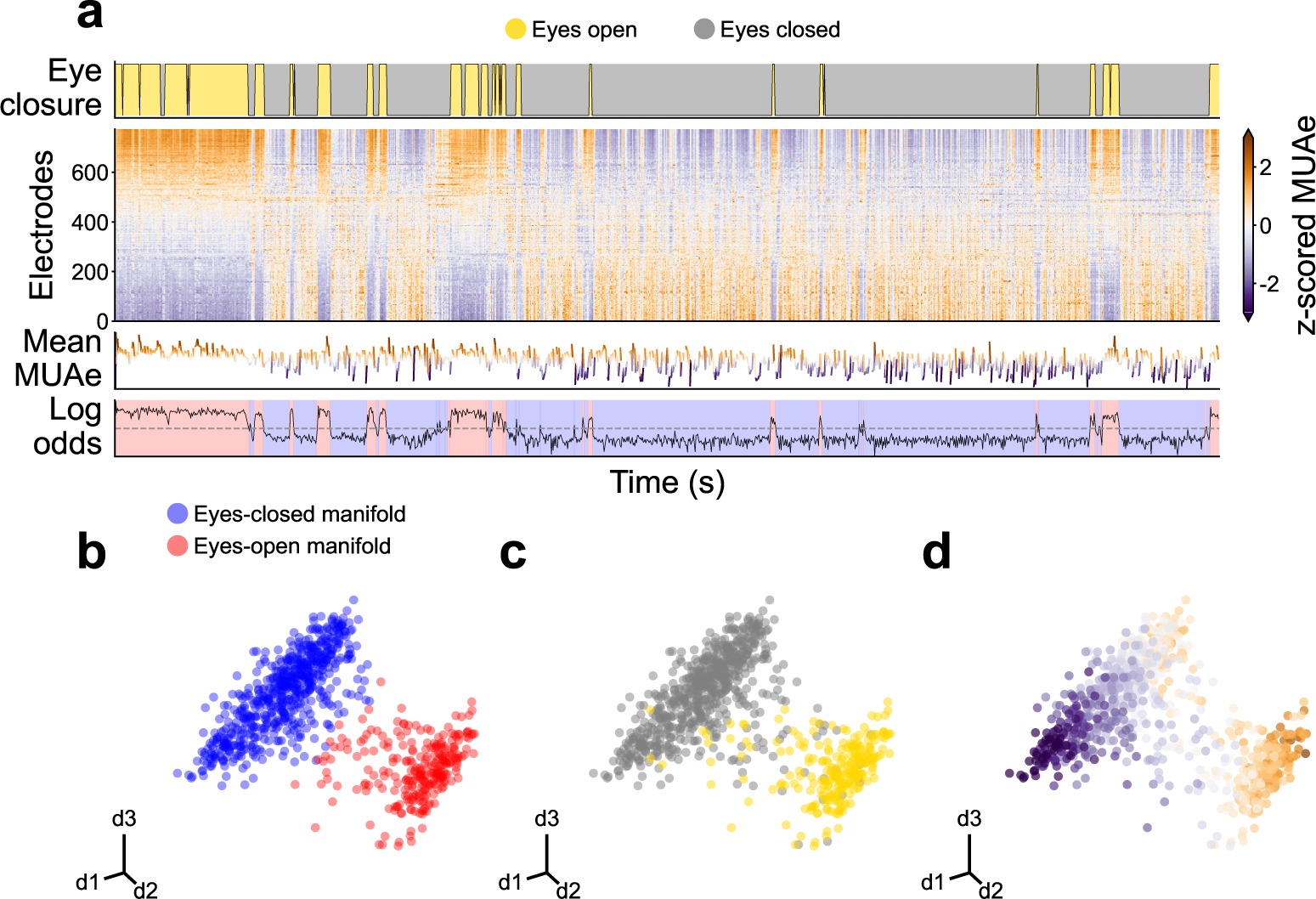
Overview of the experimental data from session L RS 100817. **a** Time evolution of signals. **b, c, d** First three principal components of the MUAe neural manifold.

**Figure S3:**
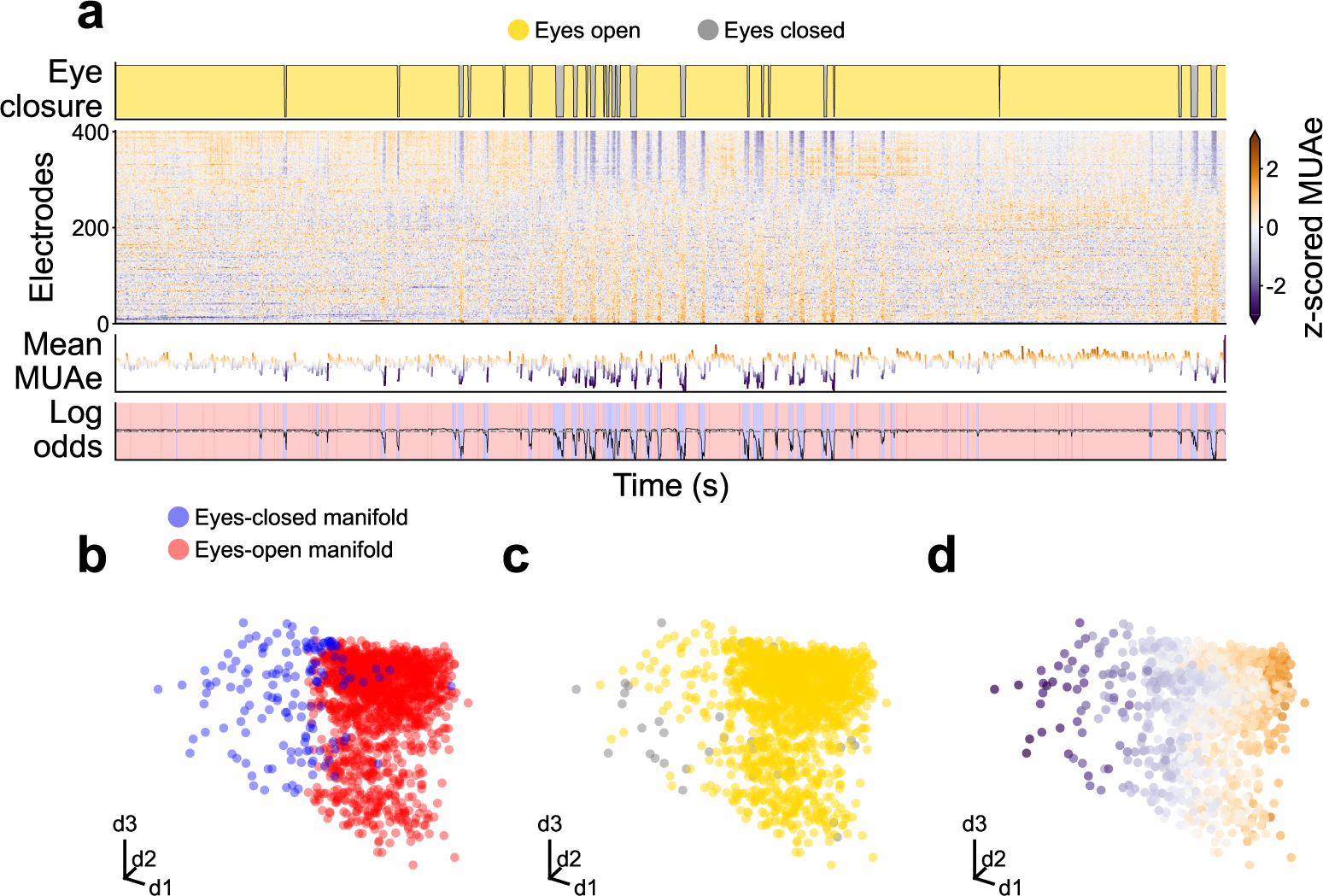
Overview of the experimental data from session A RS 150819. **a** Time evolution of signals. **b, c, d** First three principal components of the MUAe neural manifold.

**Figure S4:**
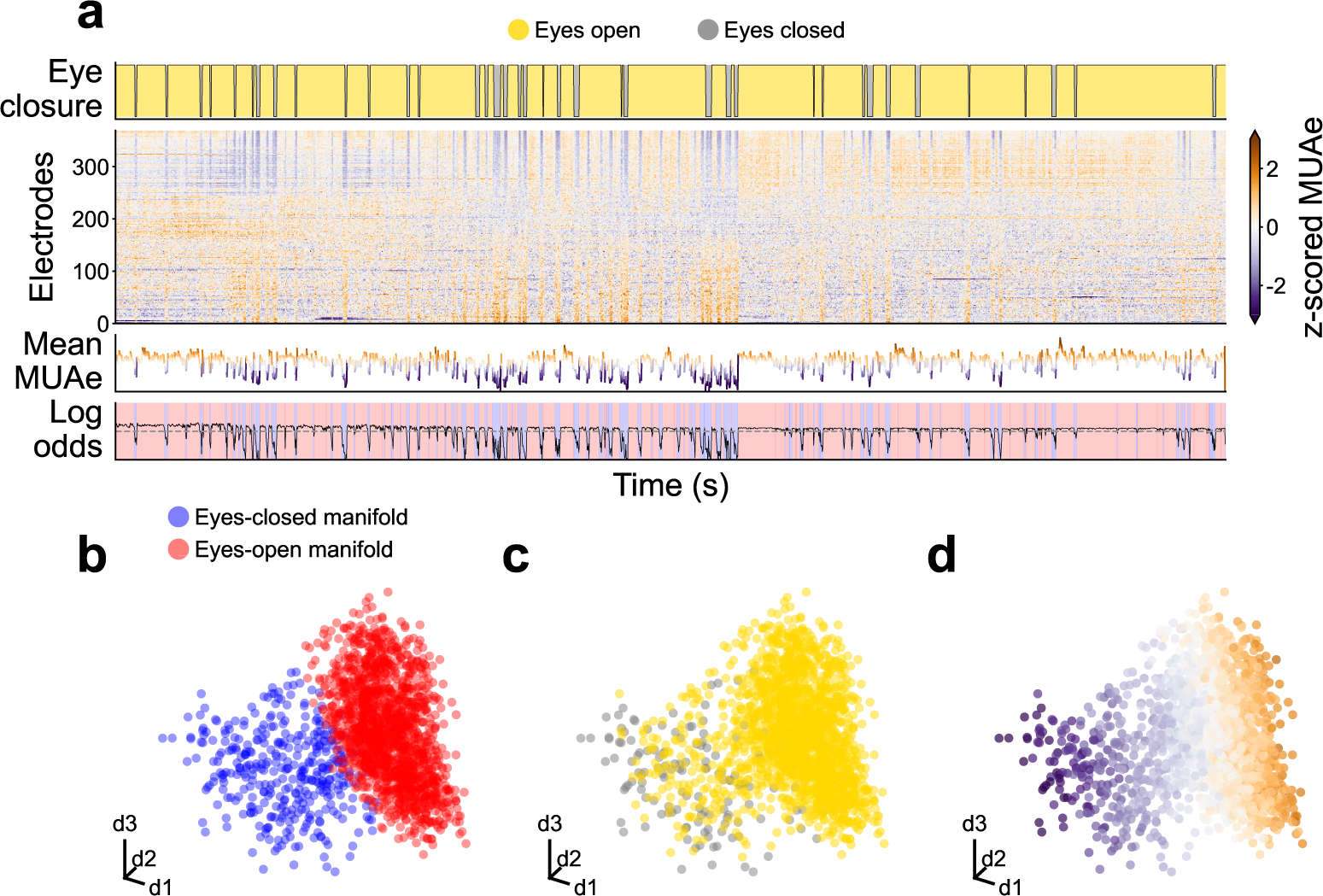
Overview of the experimental data from session A RS 160819. **a** Time evolution of signals. **b, c, d** First three principal components of the MUAe neural manifold.

**Figure S5:**
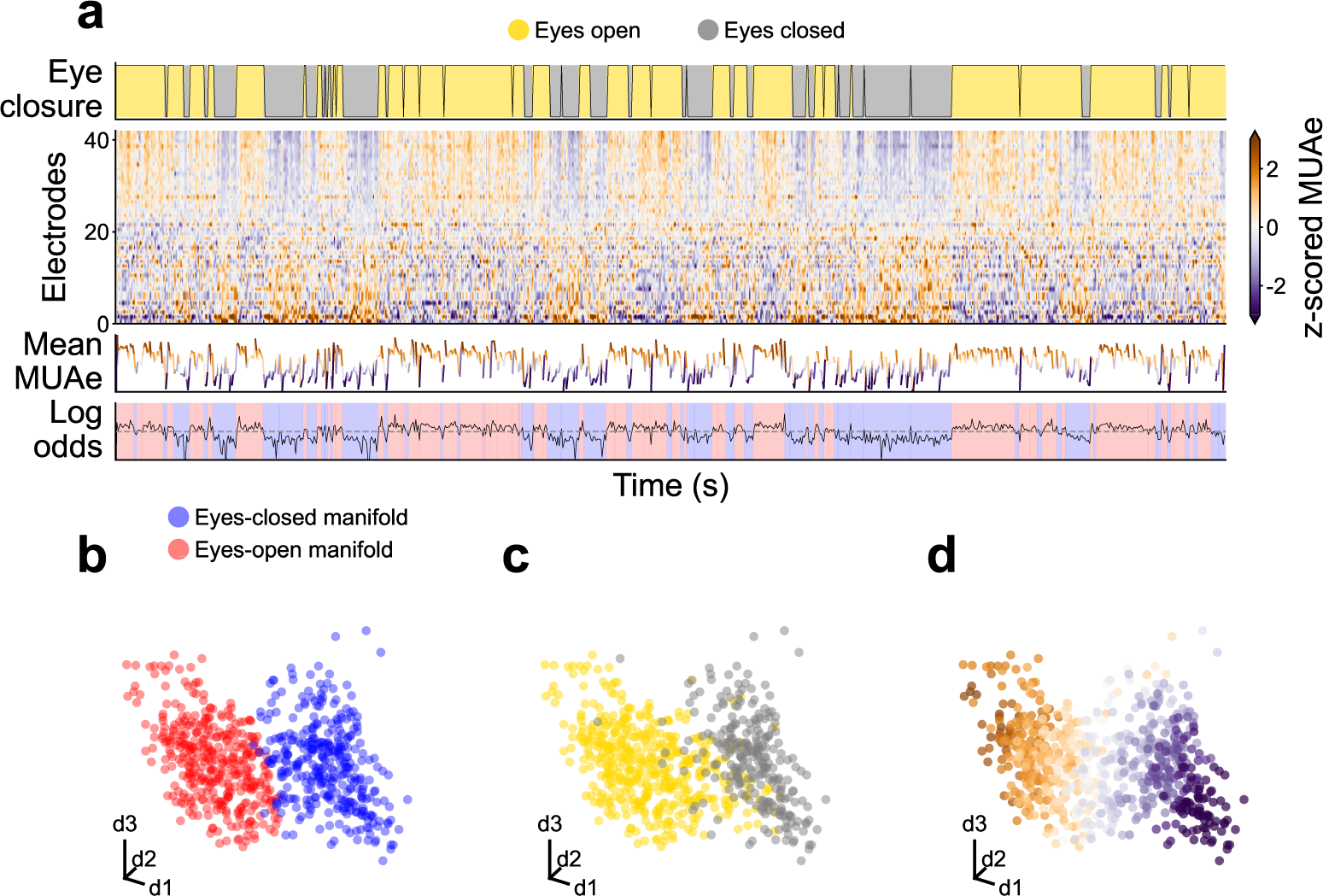
Overview of the experimental data from session Y RS 180122. **a** Time evolution of signals. **b, c, d** First three principal components of the MUAe neural manifold.

**Figure S6:**
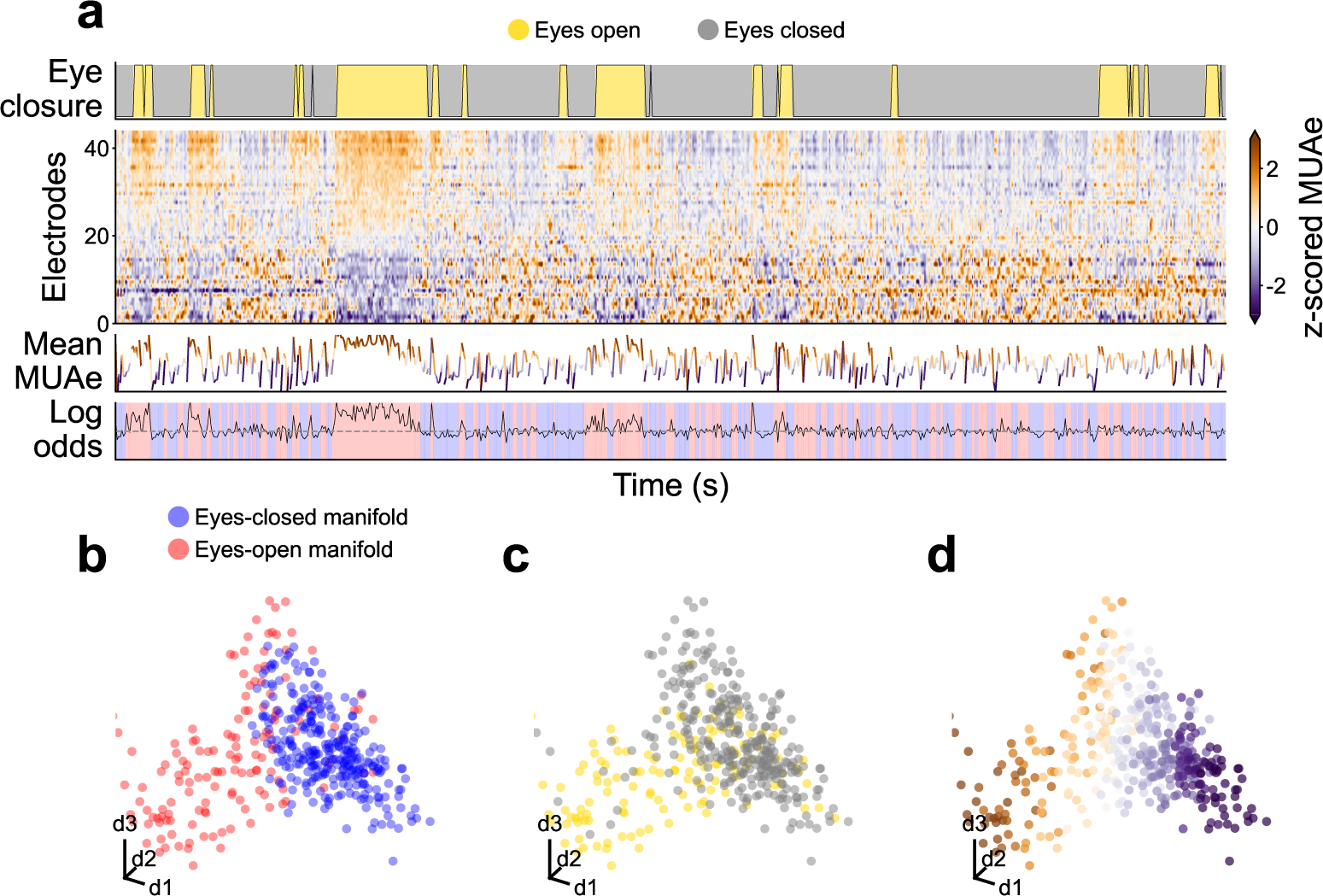
Overview of the experimental data from session Y RS 180122. **a** Time evolution of signals. **b, c, d** First three principal components of the MUAe neural manifold.

**Figure S7:**
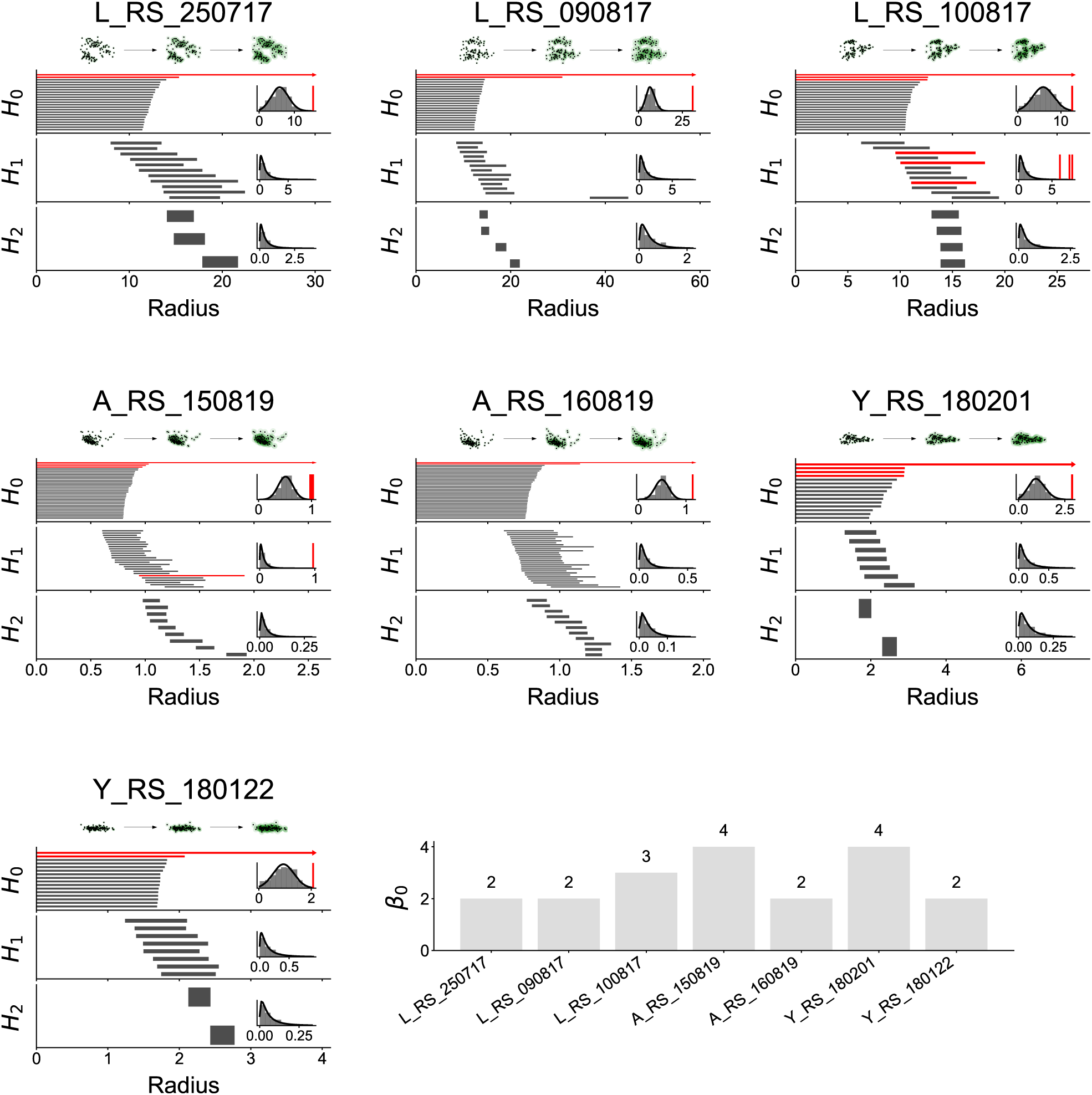
Persistence homology of the high-dimensional manifolds show the presence of at least two clusters. Each panel shows data for one session. For each panel, (Top) Sample clouds with a green radius around them. These correspond to the radius used in the persistent homology process. (Main plots) Persistence barcodes of the Vietoris-Rips complex of the 10D neural manifolds, for all sessions. (Inset plots) Distribution of barcode length with a fitted lognormal distribution, long barcodes colored red (determined ad hoc). (Bottom right panel) Number of clusters found in each session.

**Figure S8:**
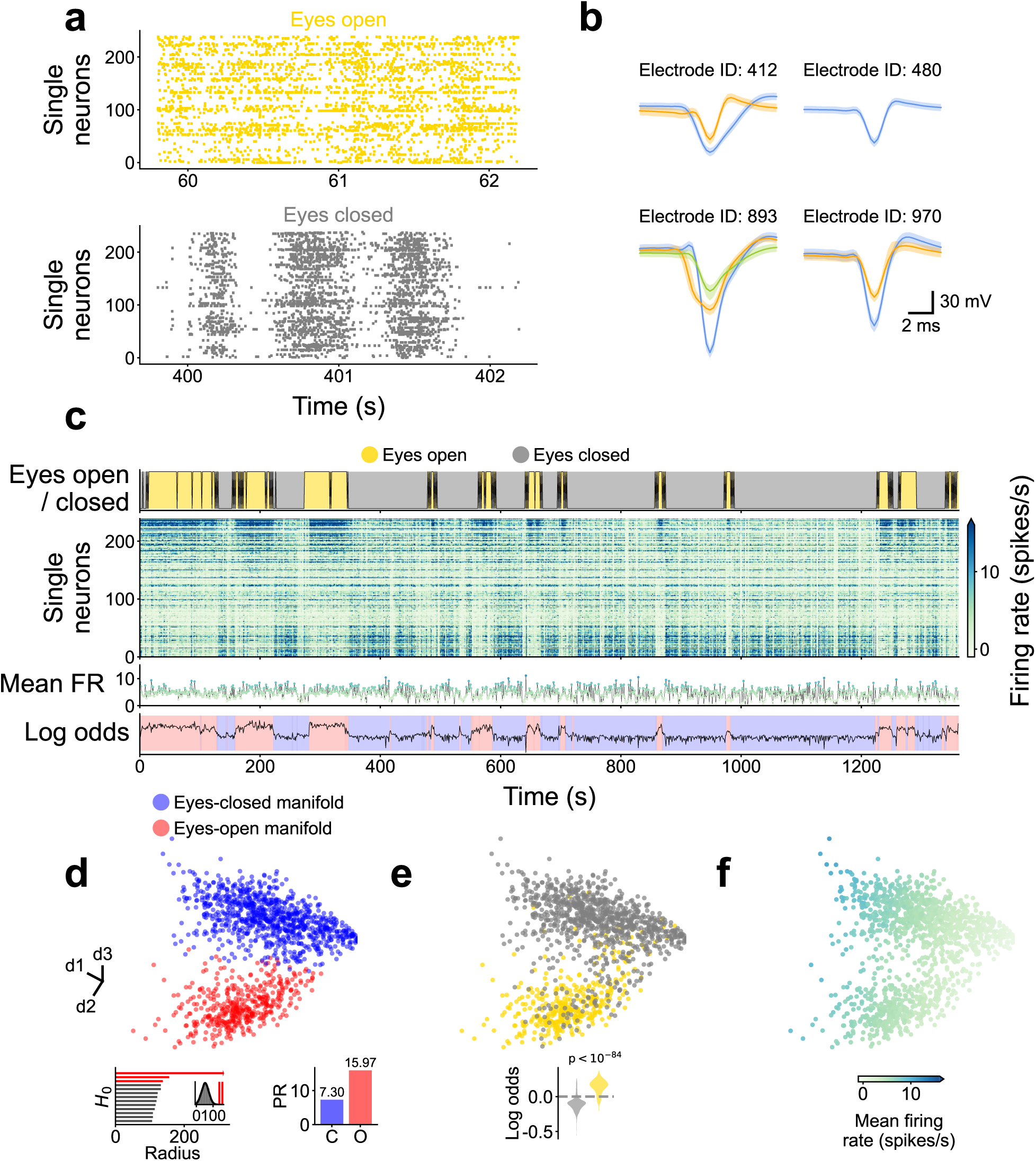
Overview of the spiking data from session L RS 250717. Single neurons were isolated using a semi-automatic spike sorting method, see Methods, Spike sorting. The firing rate (FR) was calculated counting the number of spikes in 1-second bins. a Sample spike raster plots for eyes-open and eyes-closed periods. b Sample waveforms from four electrodes, multiple single units isolated in some electrodes (color-coded). Median (solid line) and 20-80 percentiles (shading) shown per unit. c Time evolution of signals. d, e, f First three principal components of the FR. Insets show the persistent homology for the *H*_0_ homology group (d), the Participation Ratio (PR) of each manifold that are in agreement with the MUAe measurements (d) and the violin plots of the eye closure against the clustering log odds (e).

**Figure S9:**
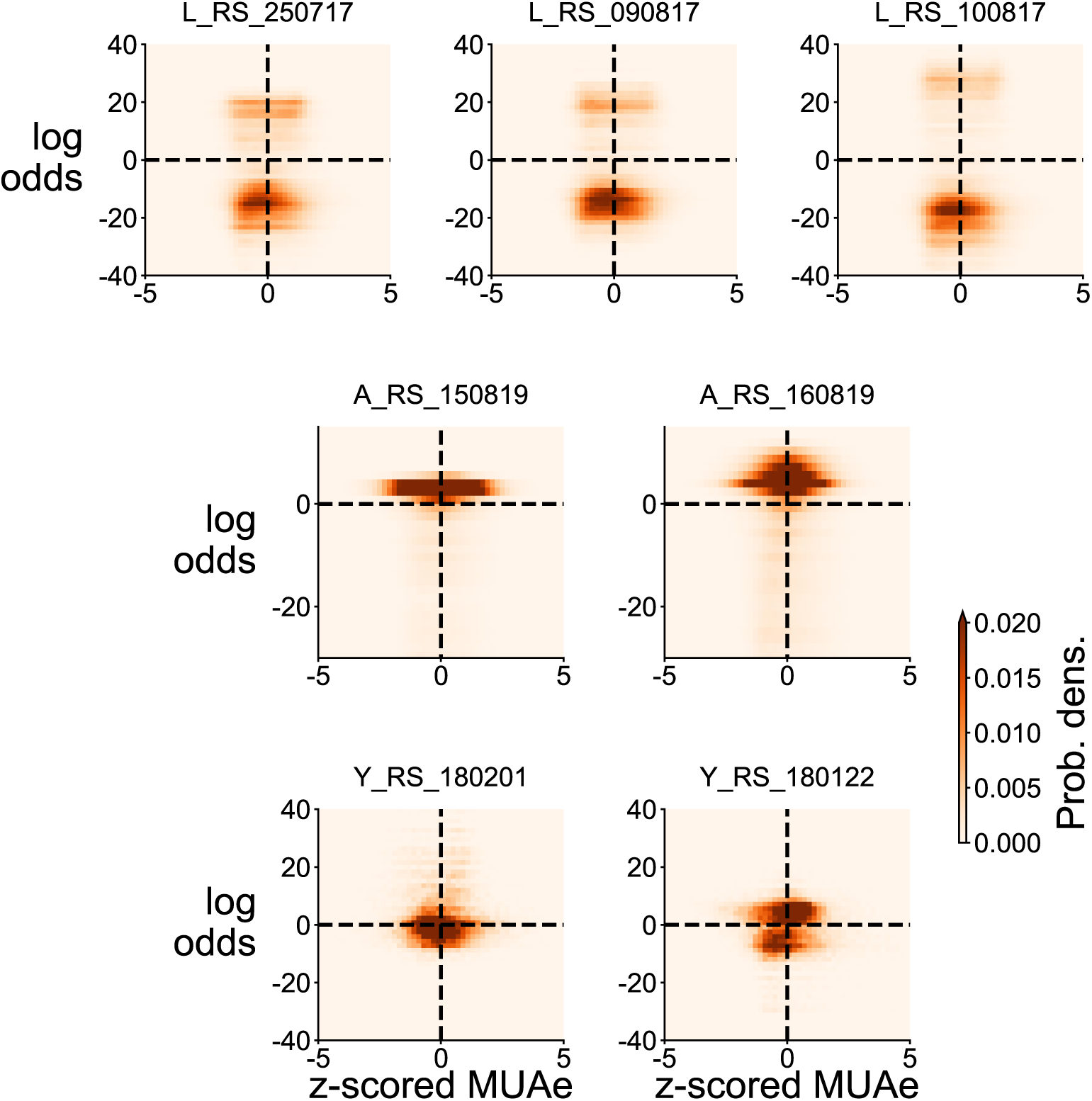
2D histograms of z-scored MUAe and log odds. Darker color indicates higher occurrence. If the neural manifolds were solely explained by the higher activity, the histograms should be strictly diagonal; we instead observe that the histograms spread across multiple quadrants and are even bimodal in some cases.

**Figure S10:**
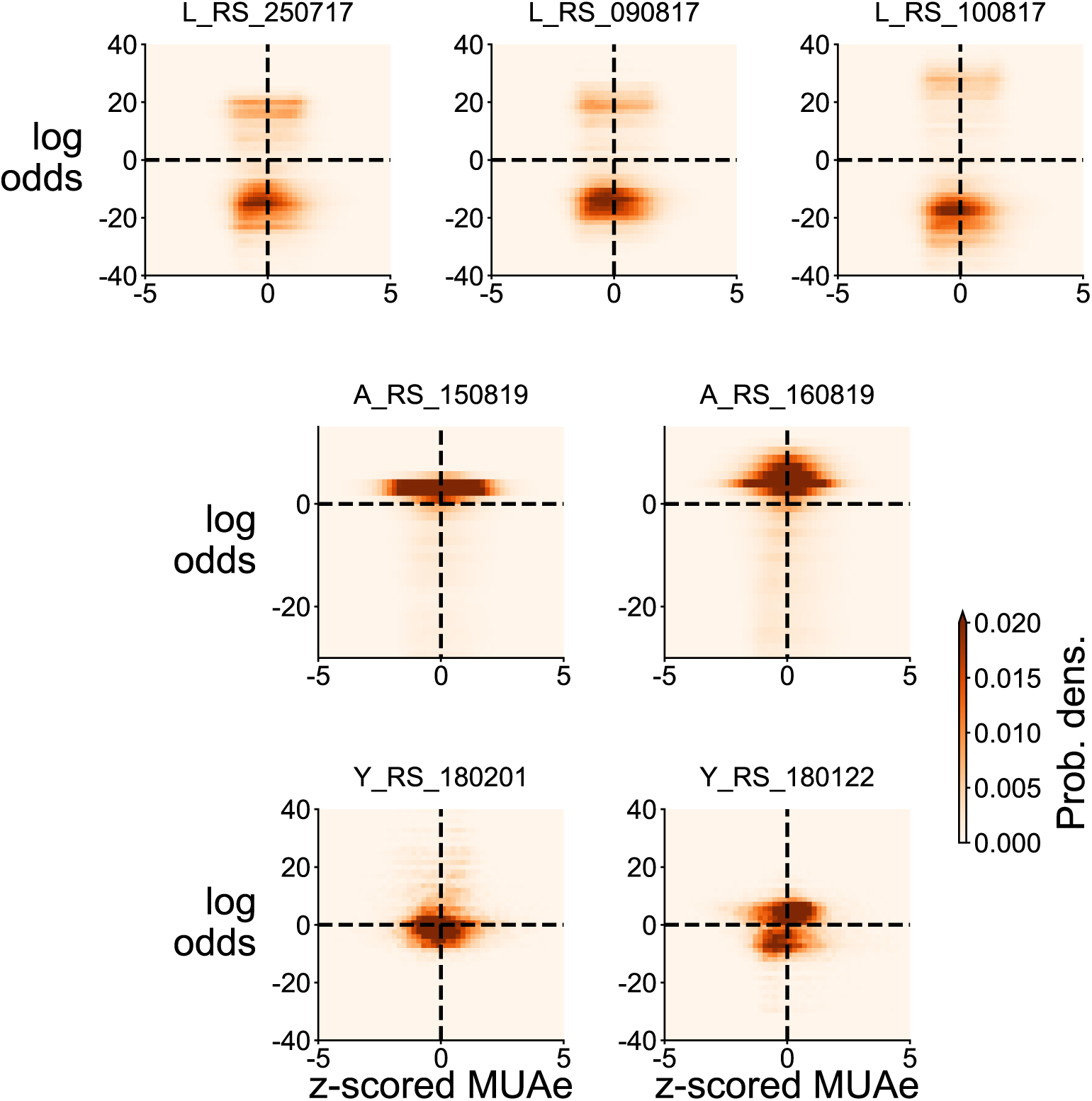
V4 activity from session L RS 250717 does not show distinct clusters in its neural manifold. **a** Time evolution of signals. **b, c, d** Three dimensional PCA of the MUAe neural manifold.

**Figure S11:**
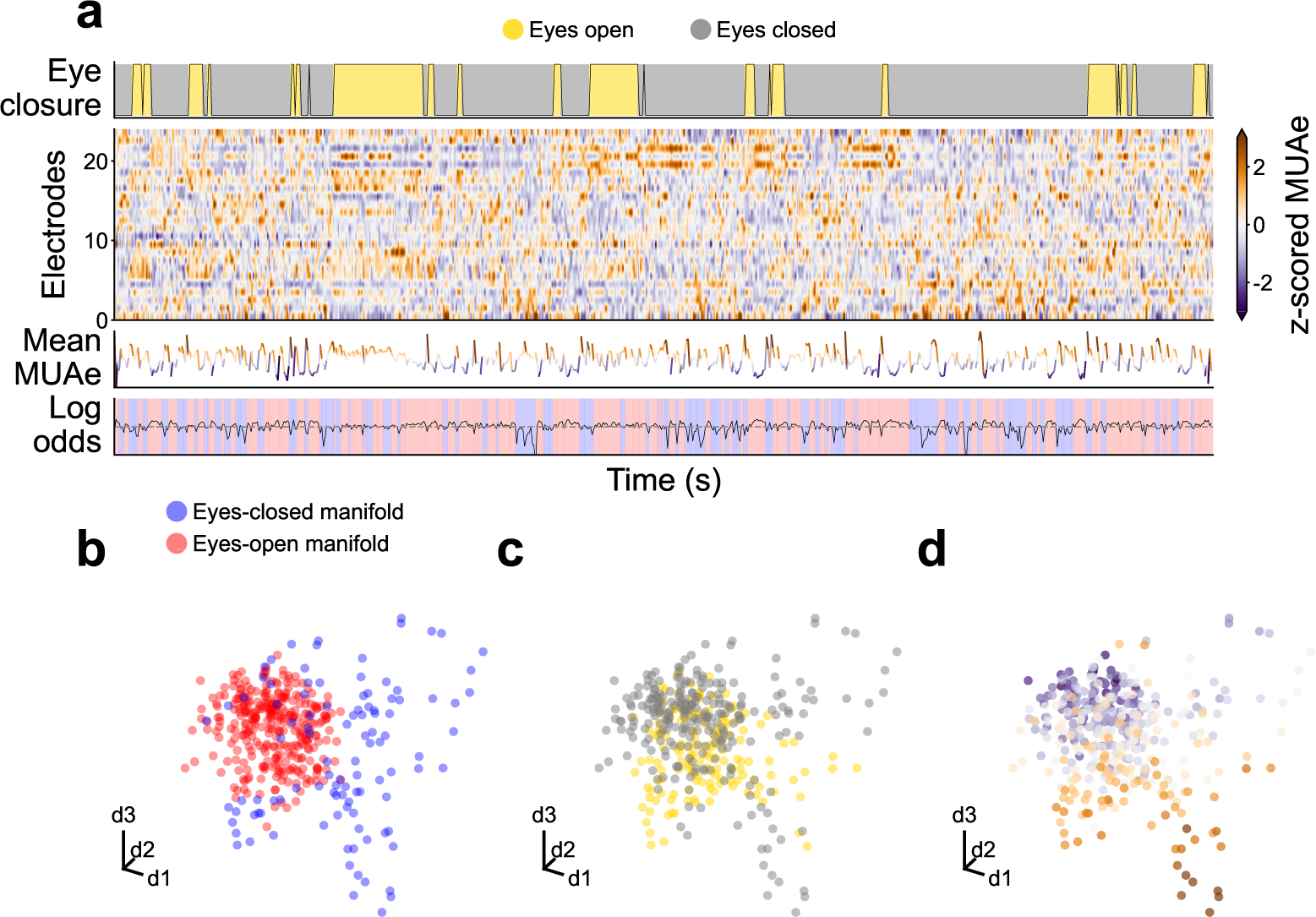
DP activity from session Y RS 180201 does not show distinct clusters in its neural manifold. **a** Time evolution of signals. **b, c, d** Three dimensional PCA of the MUAe neural manifold.

**Figure S12:**
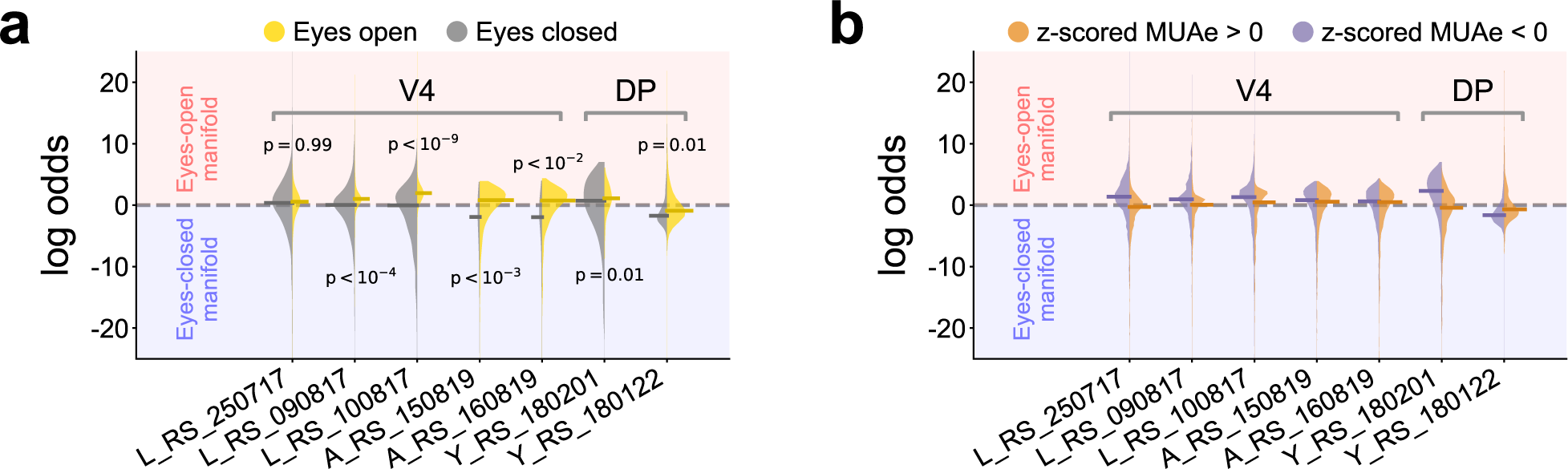
V4 and DP manifold log odds are not strongly correlated with eye closure nor with MUAe. a, b Violin plots of V4 and DP for eye closure (a) and MUAe activity (b). Neither show a clear separation along different neural manifolds.

**Figure S13:**
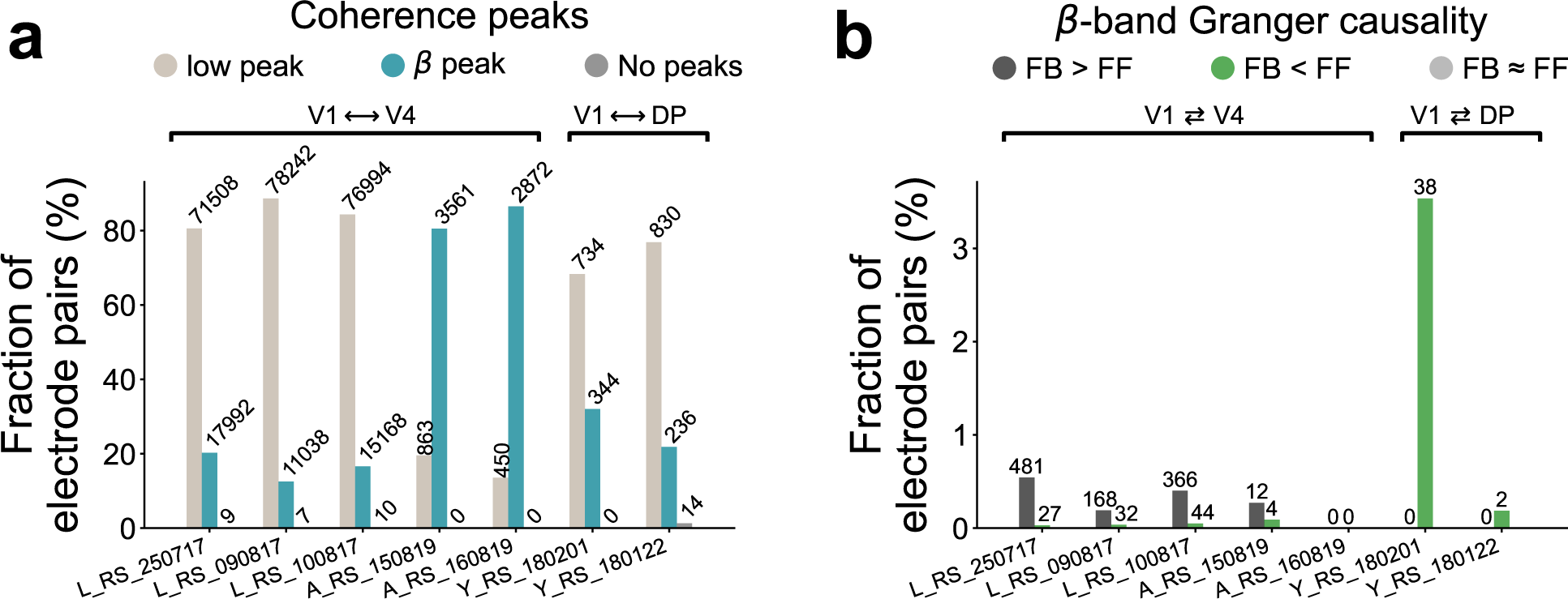
Quantification of coherence peaks and beta-band spectral Granger causality. a Quantification of coherence peaks across all sessions. A substantial portion of all electrode pairs displayed a beta peak. Note that the percentages for a session can add up to more than 100% since the same electrode pair can have both a low-frequency and a beta peak. b Quantification of beta-band spectral Granger causality for all sessions. Welch’s t-test was used to determine whether top-down Granger causality was greater than, less than, or roughly equal to bottom-up Granger causality, within the beta frequency band. The test was only applied to those electrode pairs that showed a beta coherence peak. A large portion of V1 ⇄ V4 pairs show stronger causality in the top-down direction, while V1 ⇄ DP did not appear to have prominent top-down causality compared to bottom-up causality.

**Figure S14:**
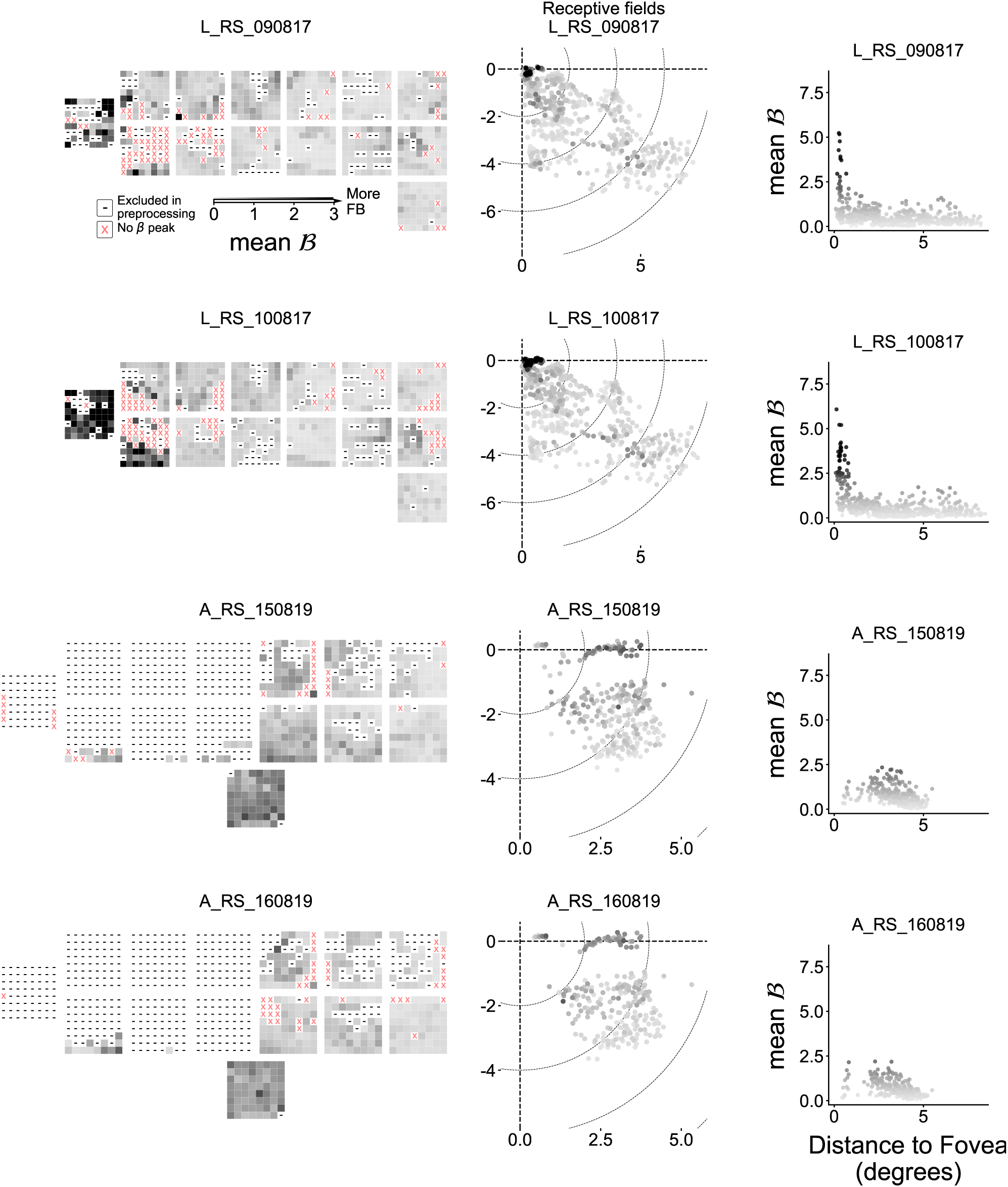
Spatial distribution of Granger causality strength per electrode for all relevant sessions. *(Left column)* Schematic representation of the electrode locations overlaid with the mean top-down signal strength B per electrode (see Coherence and Granger causality for a description of B). *(Center column)* Receptive field (RF) map overlaid with the mean B per electrode. Stronger B is found around the foveal region of V1. *(Right column)* Mean B displayed against the distance from the fovea.

**Figure S15:**
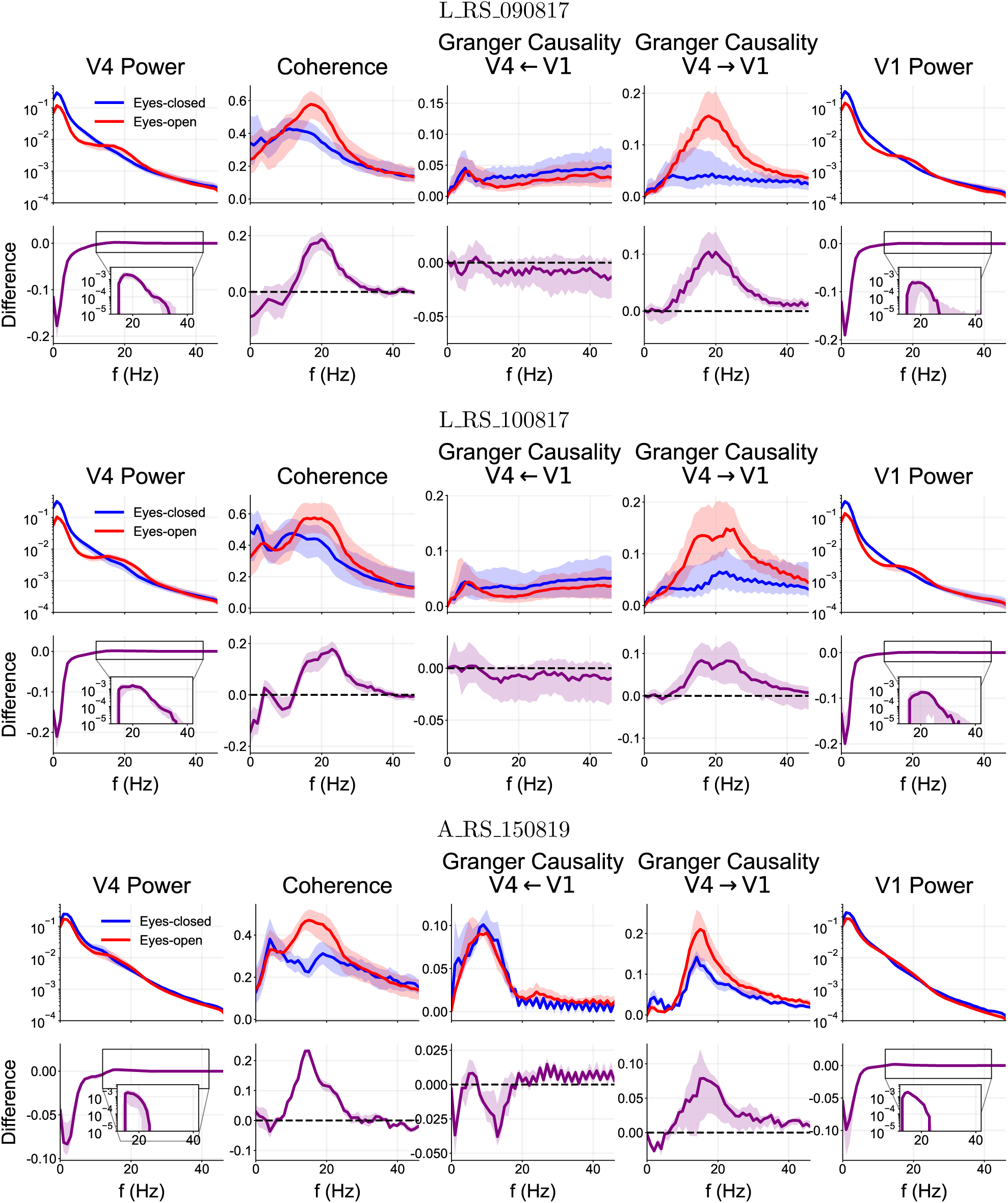
Spectral power, coherence, and Granger causality for the electrodes with high causality strength ( *>* 10) in sessions L RS 090817, L RS 100817, and A RS 150819. The data for each behavioral condition (eyes-open/closed) were concatenated and their metrics reported separately. Thick line shows median across electrodes (or pairs of electrodes) and shading indicates the 25th to 75th percentile (top row for each session). The difference between eyes-open and eyes-closed was calculated for each electrode or pair of electrodes (bottom row for each session).

**Figure S16:**
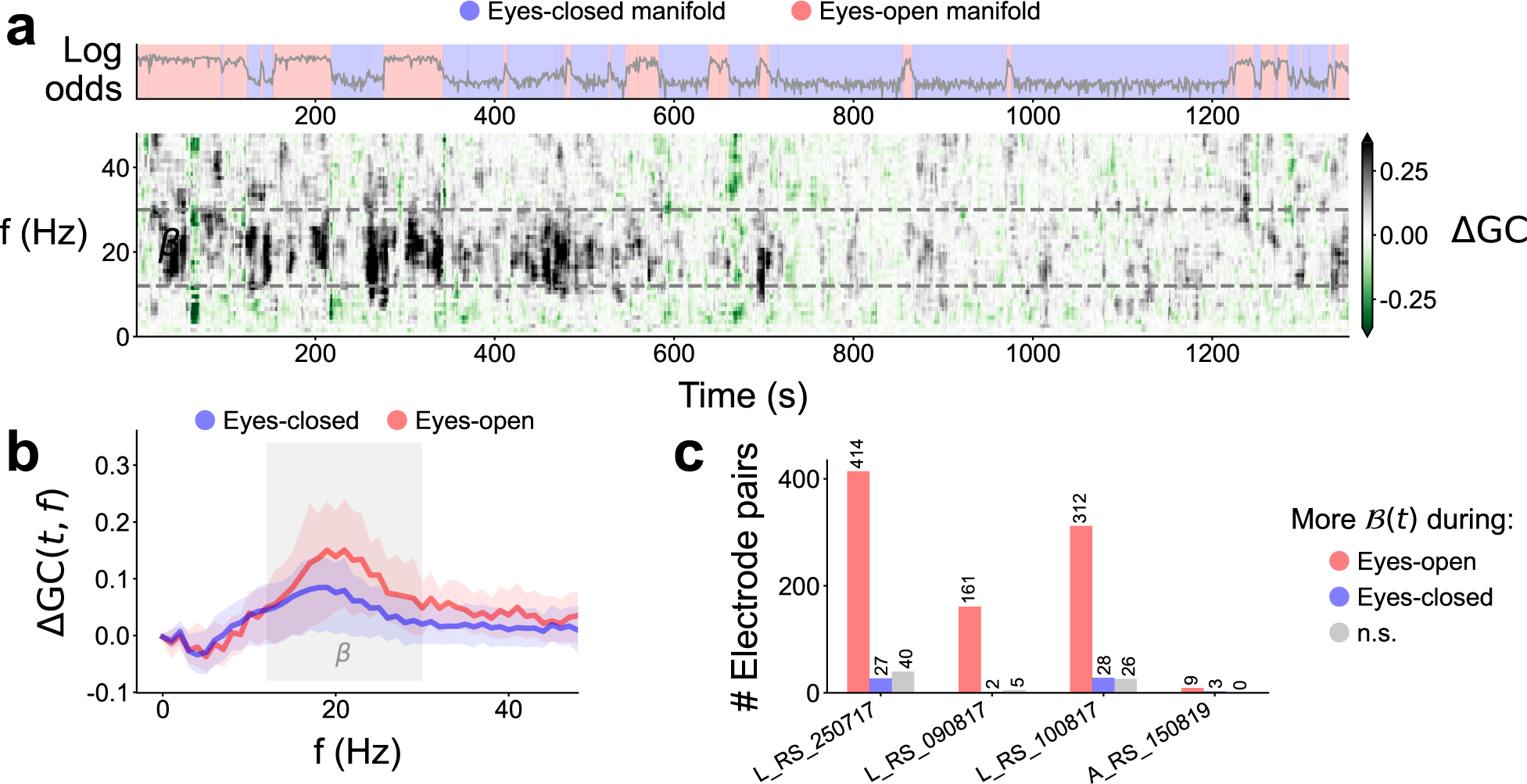
Time-dependent spectral Granger causality reveal higher top-down signals in the eyes-open periods. **a** Time evolution of the log odds (top), and the spectral Granger causality difference for a representative sample of V1-V4 electrodes (bottom). The sample electrodes were the same as in Figure 5a. **b** Causality difference median (line) and 25th to 75th percentiles (shade) in each manifold for one sample V1-V4 electrode pair. Beta frequency range highlighted. **c** Quantification of beta-band causality difference (*t*) over time (in each manifold) for all V1-V4 and V1-DP electrodes in all sessions—using Welch’s t-test.

**Figure S17:**
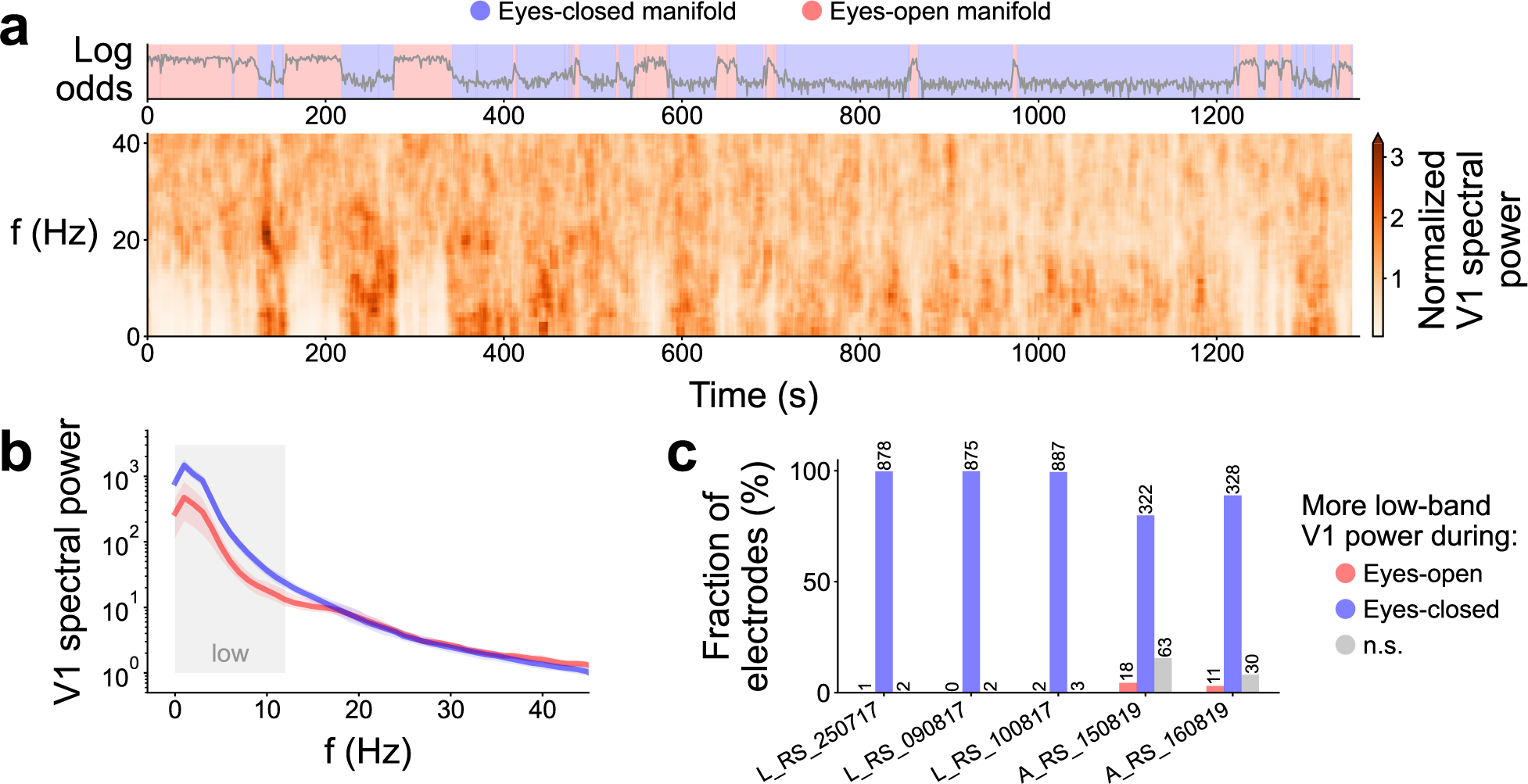
Analysis of the V1 LFP spectrogram. **a** Log odds identifying the neural manifolds (as in Figure 2a), and time-varying spectrum of a sample V1 electrode (session L RS 250717, power normalized for each frequency). **b** Spectrum of a sample V1 electrode (session L RS 250717). Colors indicate the different manifolds. **c** Result of t-test in the low frequency band (less than 12 Hz) for all V1 electrodes. As expected, the overwhelming majority of electrodes displays higher low frequency power when the eyes are closed.

**Figure S18:**
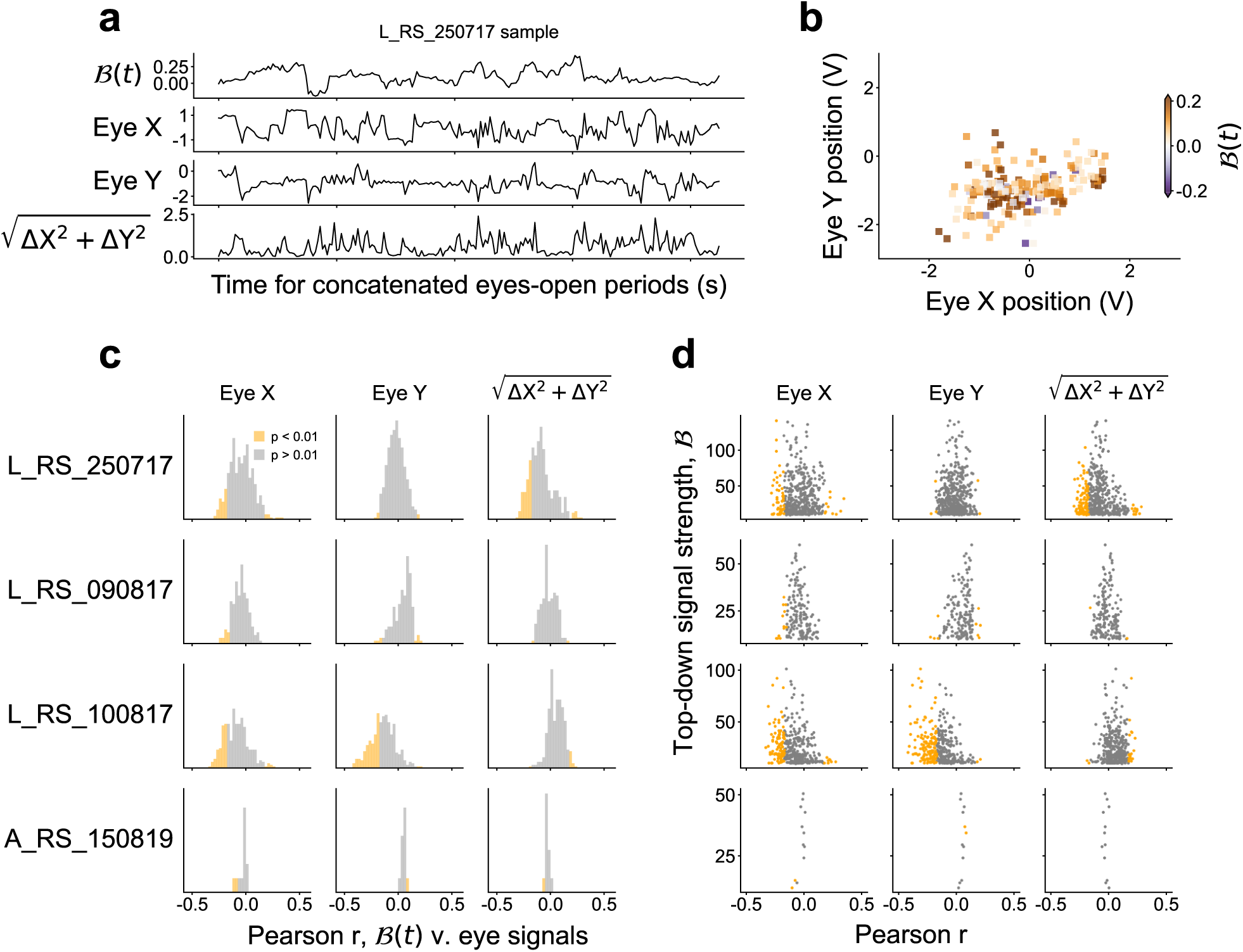
Top-down signals are not correlated with gaze direction. **a** Sample t_√_ra<u>ces of the b</u>eta-band Granger causality difference, gaze direction (Eye X, Eye Y), and gaze movement speed 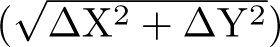. **b** Sample beta causality difference over the gaze locations. Higher top-down causality is not concentrated in particular regions. **c** Histograms of Pearson correlation coefficients between time-dependent causality difference and gaze signals, computed for all electrode pairs in all sessions. Significant (*p <* 0.01 two-tailed) part of histograms shown in orange. Note that we did not correct for multiple testing, since reducing the p-value threshold would simply reinforce our finding that no strong correlation was present between the gaze and the top-down signals; i.e., no multiple testing correction was the more conservative approach in this case. **d** Scatter plot of the summed time-independent causality difference against the correlation with gaze direction and movement. There is no clear relation between B(*t*)-gaze correlation and causality strength.

**Figure S19:**
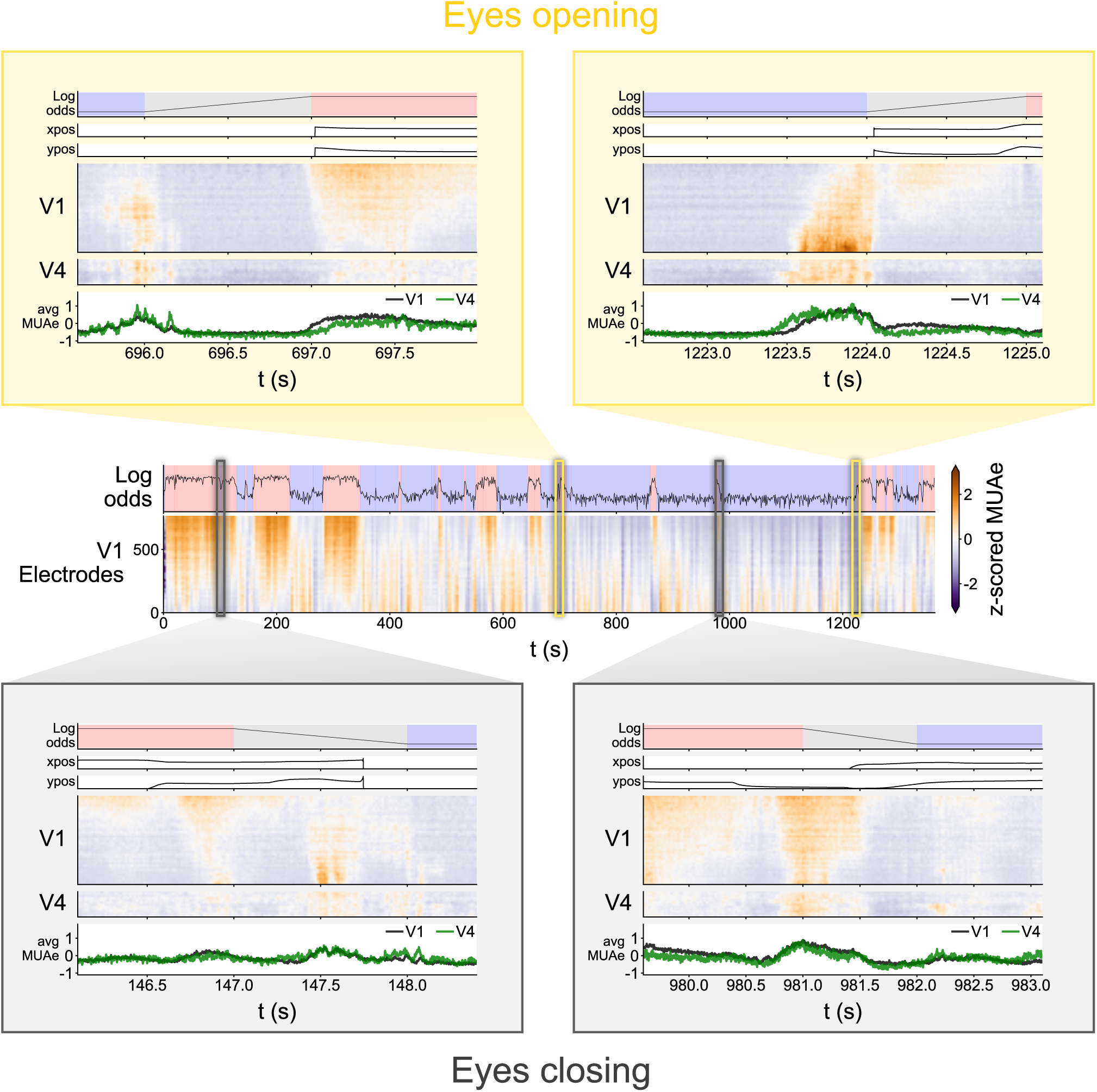
Closer look at the V1 and V4 MUAe around the transitions between the two manifolds in session L RS 250717.

## Footnotes

1 spyking-circus.readthedocs.io

2 ripser.scikit-tda.org

## Notes

### Competing Interest Statement

The authors have declared no competing interest.

### Summary of Updates

Updated author affiliations due to institution renaming. Extension of discussion with further insights.

